# Engineered niches support the development of human dendritic cells in humanized mice

**DOI:** 10.1101/835223

**Authors:** Giorgio Anselmi, Kristine Vaivode, Charles-Antoine Dutertre, Pierre Bourdely, Yoann Missolo-Koussou, Evan Newell, Oliver Hickman, Kristie Wood, Alka Saxena, Julie Helft, Florent Ginhoux, Pierre Guermonprez

## Abstract

Classical dendritic cells (cDCs) are rare sentinel cells specialized in the regulation of adaptive immunity. Modeling cDC development is both crucial to study cDCs and harness their potential in immunotherapy. Here we addressed whether cDCs could differentiate in response to trophic cues delivered by mesenchymal components of the hematopoietic niche where they physiologically develop and maintain. We found that expression of the membrane bound form of human FLT3L and SCF together with CXCL12 in a bone marrow mesenchymal stromal cell line is sufficient to induce the contact-dependent specification of both type 1 and type 2 cDCs from CD34^+^ hematopoietic stem and progenitor cells (HSPCs). Engraftment of these engineered mesenchymal stromal cells (eMSCs) together with CD34^+^ HSPCs creates an *in vivo* synthetic niche in the dermis of immunodeficient mice. Cell-to-cell contact between HSPCs and stromal cells within these organoids drive the local specification of cDCs and CD123^+^AXL^+^CD327^+^ pre/AS-DCs. cDCs generated *in vivo* display higher levels of resemblance with human blood cDCs unattained by *in vitro* generated subsets. Altogether, eMSCs provide a novel and unique platform recapitulating the full spectrum of cDC subsets enabling their functional characterization *in vivo*.

## Introduction

Classical human dendritic cells (cDCs) are sentinels of the immune system with a unique ability to regulate the function of T lymphocytes ^1^. Dendritic cells can induce immune tolerance ^2^ or drive the development of immunity ^3^.

The analysis of blood circulating subsets has revealed that cDCs consist in two major subtypes: CD141^+^XCR1^+^Clec9A^+^ DCs (cDC1) and CD1c^+^CD11c^+^CD172a (SIRPα) ^+^ DCs (cDC2s) ^4–6^. Both cDC1s and cDC2s arise from a bone marrow committed progenitor ^7^ or from early IRF8^+^ multipotent progenitors ^8, 9^, which generate a common circulating precursor ^10^ that progressively diverge in pre-cDC1s and pre-cDC2s ^10–12^. Type 1 DCs are conserved between mouse and human, and they share the expression of specific surface markers such as Clec9A ^13^ and XCR1 ^5^ as well as the transcription factor (TF) IRF8, which is essential for the development of murine cDC1s ^4–6, 13–15^. Conversely, human CD1c^+^ type 2 DCs have been shown to express the IRF4 TF ^16^, which controls the development of their phenotypic equivalent in the mouse model ^16, 17^. This rather simple picture is complicated by the diversity of CD1c^+^ cells, which encompass migratory DCs as well as CD14^int^ inflammatory DCs recruited during inflammation ^18–20^. More recently, un-biased approaches have uncovered a deeper complexity in the DC network with the identification of 2 types of cDC2s with distinct transcriptional profiles and the identification of AXL^+^CD11c^+^CD1c^+^ cells which have been proposed to act as a precursor for cDCs ^12, 21^.

Human hematopoietic progenitors reside in the stem cells niche of the bone marrow. Genetic studies in the murine model identified three essential factors supporting HSPCs homeostasis: the membrane-bound form of stem cell factor (SCF/KITL) ^22, 23^, the C-X-C motive chemokine 12 (CXCL12) ^24, 25^ and thrombopoietin (TPO) ^26, 27^. In the bone marrow, perivascular mesenchymal stromal cells have been described as the main source of SCF and other niche factors ^28^. At steady state, Flt3-ligand (FLT3L) is delivered as a membrane-bound precursor expressed on radio-resistant stromal cells ^29–31^. After egressing from the bone marrow, DC precursors circulate in the blood and seed the peripheral tissues ^32^. In the lymph node, stromal fibroblastic reticular cells provide FLT3L ^33^, and FLT3-dependent proliferation of cDC in periphery is required for their maintenance ^34, 35^.

Modelling the development of cDCs in culture systems is instrumental to better understand their ontogeny and define their immunological function. Pioneer work from Banchereau *et al*. have identified that GMCSF and TNF-α cooperate to produce CD1a^+^ cells with features of Langerhans cells from CD34^+^ hematopoietic stem and progenitor cells (HSPCs) ^36^. Sallusto *et al*. have shown that GMCSF and IL-4 induce the differentiation of CD1c^+^CD1a^+^ inflammatory DCs from CD14 monocytes ^37^. More recent work has demonstrated that FLT3L (with TPO or with SCF/KITL, GM-CSF and IL-4) is instrumental in generating CD141^+^ cDC1s aligning phenotypically and functionally with cDC1s ^7, 8, 38–40^. This is in line with the crucial role of FLT3L, engaging the Flt3 receptor tyrosine kinase ^41–43^ in controlling DC homeostasis both in mice ^32, 34, 44, 45^ and humans ^10, 29, 46, 47^. Moreover, the activation of Notch signaling pathway has been shown to further improve the *in vitro* differentiation of both human and mouse cDC1s ^48, 49^. Despite the successes in modeling cDC1 differentiation *in vitro*, CD1c^+^ cells found in culture of CD34^+^ HSPCs either align poorly with *bona fide* blood circulating cDC2s ^39^ or were not extensively characterized ^48, 49^.

Recapitulating human cDC development *in vivo* has the potential to greatly improve our understanding of DC biology and facilitate its translational applications. Human cDCs have been found in stably reconstituted humanized mice treated with supraphysiological concentration of human FLT3L ^29, 50, 51^. However, the generated CD11c^+^CD141^+^ and CD11c^+^CD1c^+^ cells were poorly characterized and their dissemination to peripheral tissues has rarely been assessed ^52^.

Here we aimed at modeling human cDC development by providing physiological factors associated to hematopoietic niches. We found that engineered mesenchymal stromal cells (eMSCs) expressing a combination of membrane-bound FLT3L and SCF/KITL together with CXCL12 provide a scaffold for human cDC differentiation. Engraftment of eMSCs along with CD34^+^ HSPCs leads to the local development of cDCs in immunodeficient mice. This *in vivo* system recapitulates the differentiation of pre/AS-DCs, cDC1s and cDC2s with an unreached level of similarity with the phenotype of human blood cDCs.

## Results

### Stromal membrane-bound FLT3L is sufficient to support human cDC differentiation from CD34^+^ HSPCs

We hypothesized that the interaction of hematopoietic progenitors with membrane bound factors expressed by stromal cells of the niche would promote the specification of the cDC lineage.

To test this, we engineered a bone marrow-derived murine mesenchymal cell line (MS5) ^7, 53^ to stably and homogeneously express the transmembrane form of human FLT3L (MS5_F) as probed by flow cytometry (Fig. 1a). Co-culture of MS5_F with CD34^+^ HSPCs drives the appearance of cDC1-like Clec9A^+^CD141^+^ and cDC2-like CD14^−^CD1c^+^ cells. Importantly, MS5_F is more efficient than recombinant soluble FLT3L (MS5+recFL) in generating cDC-like cells (Fig. 1b).

**Figure 1.**
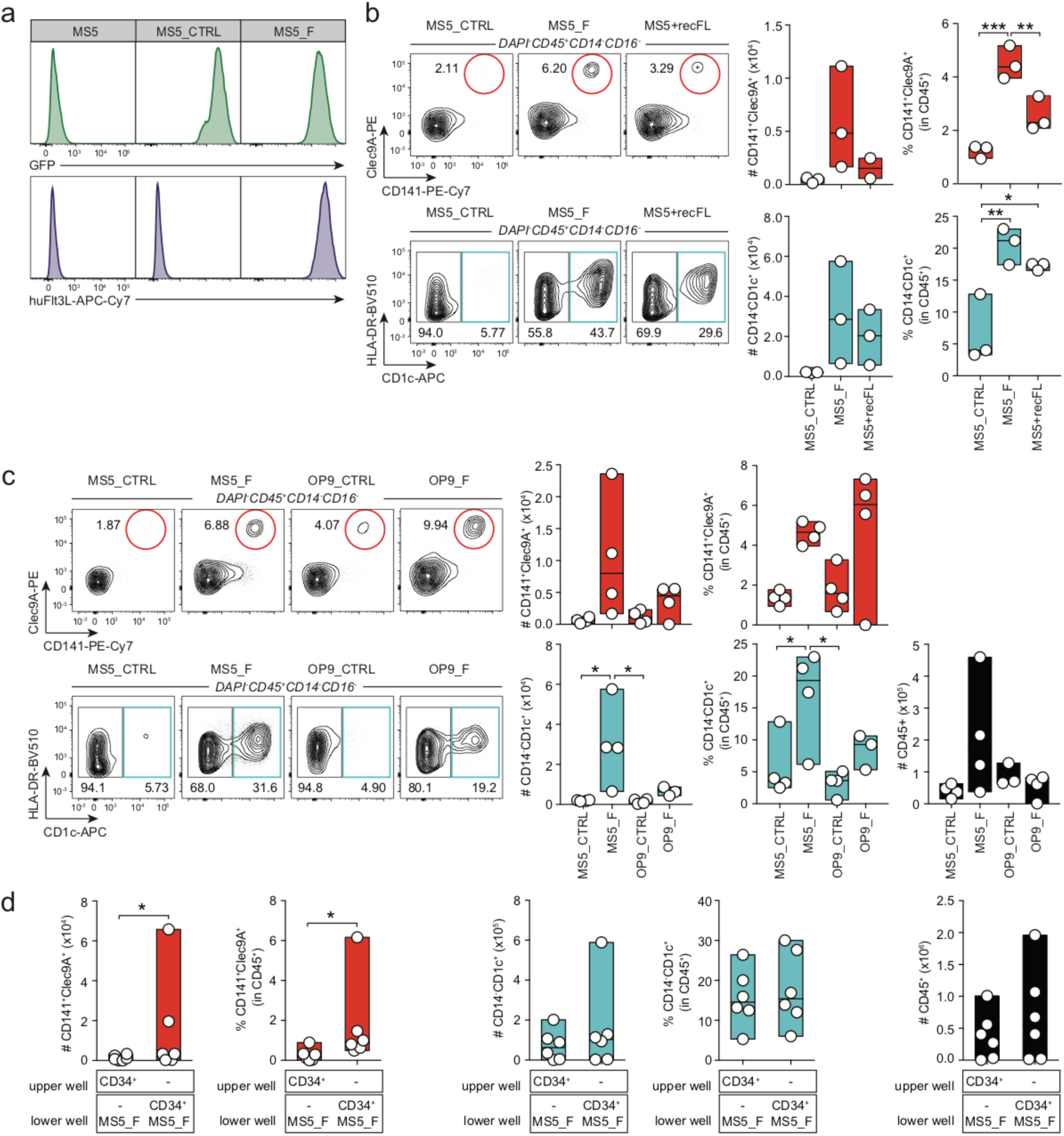
Stromal membrane bound FLT3L efficiently supports human DC differentiation from CD34^+^ HSPCs. (a) Expression of membrane bound FLT3L in mouse bone marrow-derived stromal cells engineered to express human FLT3L (MS5_F) and control (MS5_CTRL). (b) Human cDC subsets differentiated *in vitro* from CD34^+^ cord blood-derived HSPCs cultured with MS5 expressing membrane bound FLT3L (MS5_F) or MS5 supplemented with recombinant human FLT3L (MS5+recFL) at day 15 (n=3 donors in one experiment. Line represents median; * p<0.05, ** p<0.01, *** p<0.001, one-way ANOVA test). (c) Representative flow cytometry plots and quantification of human cDC subsets differentiated *in vitro* from cord blood-derived CD34^+^ progenitors in culture with mouse stromal cell lines MS5 and OP9 expressing human FLT3L (MS5_F and OP9_F) at day 15 (n=4 donors in one experiment. Line represents median; * p<0.05, one-way ANOVA test). (d) Absolute number and frequency of CD141^+^Clec9A^+^ and CD14^−^CD1c+ human cDCs differentiated from CD34^+^ HSPCs in direct contact (lower well) or physically separated (upper well) from engineered MS5_F. DC differentiation was assessed at day 15 by flow cytometry (n=6 donors in 3 independent experiments. Line represents median; * p<0.05, two-tailed paired Student t test).

In contrast, OP9 ^54^ hematogenic stromal cells stably expressing membrane bound FLT3L (OP9_F) were less efficient than MS5_F in driving cDC differentiation (Fig. 1c). Besides, MS5_F also promoted the appearance of CD123^+^CD303/4^+^ cells resembling either pDCs or pre/AS-DCs ^12, 21^ (Supplementary Fig.1a and b).

Next, we wanted to test whether cell-to-cell interactions mediate the differentiation of cDCs driven by FLT3L-expressing MS5 stromal cells. Using transwell permeable to soluble factors but preventing cognate interactions, we found that direct contact is required to support efficiently cDC differentiation (Fig. 1d).

Collectively, these data show that membrane FLT3L expression in stromal cells provide an improved platform to trigger the differentiation of cDC-like cells from CD34^+^ HSPCs *in vitro via* cell-to-cell contact.

### Stromal CXCL12 and membrane bound SCF improve FLT3L-driven development of human cDC *in vitro*

Next, we sought to improve the efficiency of cDC production in MS5_F by co-expressing additional niche factors. We focused on SCF, CXCL12 and TPO because of their essential role in supporting HSPCs maintenance in the bone marrow niche ^22, 24–27, 55^. SCF had also been extensively used in previously published DC culture protocols ^7, 39, 40, 56^.

To this end, we generated a collection of MS5 stromal cells stably expressing either one, two, three or four human factors by combining CXCL12, TPO and membrane-bound SCF/KITL, with or without membrane-bound FLT3L (Supplementary Fig. 2a).

We screened this collection of engineered mesenchymal stromal cell (eMSC) lines based on their ability to support human cDC differentiation from cord blood-derived CD34^+^ HSPCs.

At day 15, only FLT3L-expressing eMSCs successfully supported the differentiation of CD141^+^Clec9A^+^ and CD14^−^CD1c^+^ cells (Fig. 2a and Supplementary Fig. 3a). We conclude that FLT3L is necessary for the differentiation of cDCs using eMSCs. Besides, optimal cDC production was obtained in cultures containing eMSC co-expressing membrane bound SCF and CXCL12 together with FLT3L (MS5_FS12) (Fig. 2a), whereas no difference was observed for CD14^+^CD16^−^ monocytes and CD14^+^CD16^+^ macrophages as compared to MS5_CTRL (Supplementary Fig. 3b).

**Figure 2.**
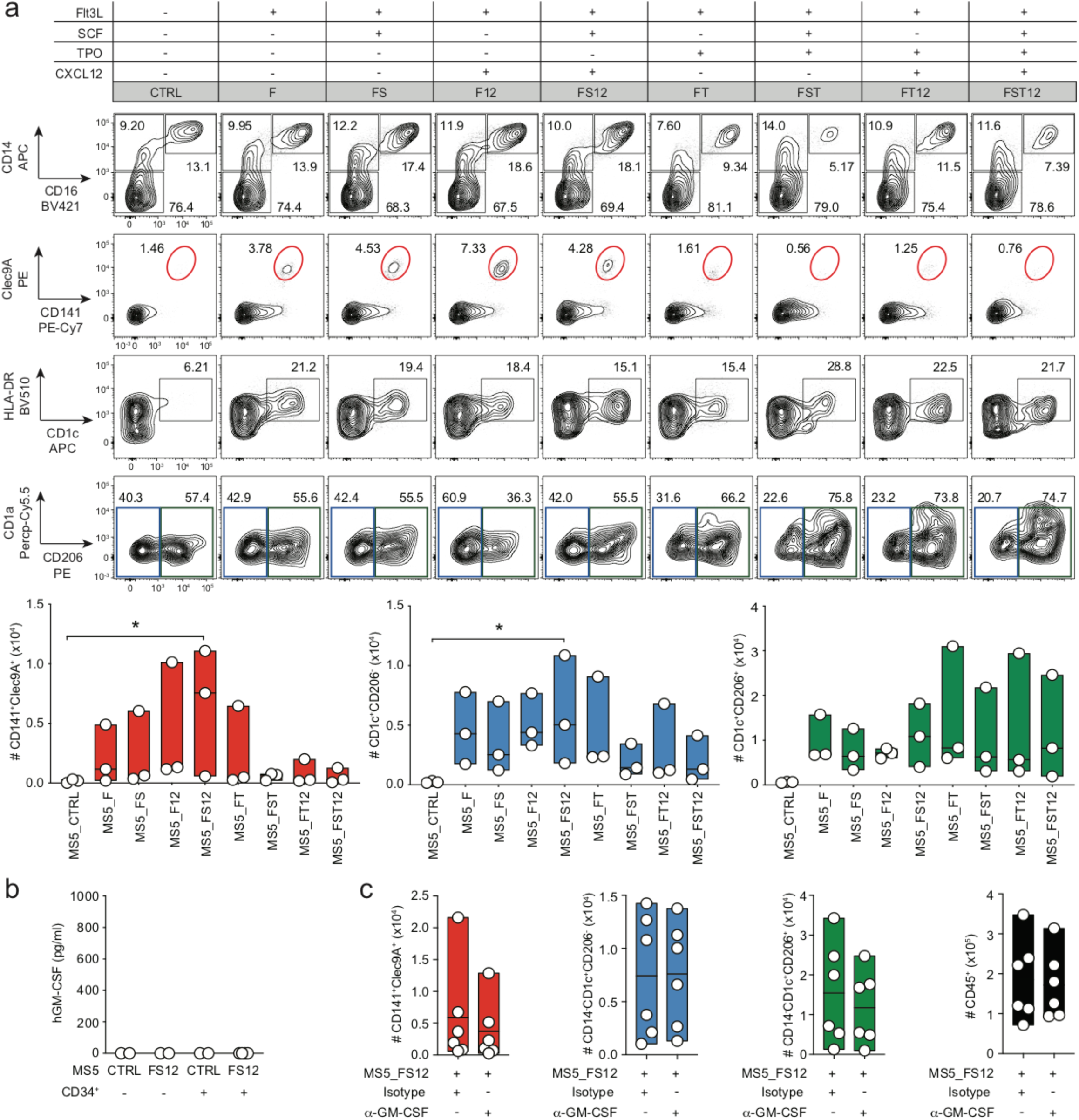
Stromal membrane bound SCF and CXCL12 improve the FLT3L-driven development of human cDC *in vitro*. (a) Representative FACS plots and absolute number of CD141^+^Clec9A^+^, CD1c^+^CD206^−^ and CD1c^+^CD206^+^ cells generated from CD34^+^ HSPCs cultured with MS5 expressing human FLT3L (MS5_F) in combination with human SCF (S), TPO (T) and CXCL12 (12). Day15 flow cytometry analysis of n=3 cord blood donors in 3 independent experiments (line represents median; * p<0.05, one-way ANOVA test). (b) ELISA detecting human GM-CSF in the supernatant of CD34^+^ HSPCs cultured with engineered MS5 expressing human FLT3L, SCF and CXCL12 (MS5_FS12) at day 15 (mean ± SEM of n=2-4 independent experiments). (c) Absolute number of human DC subsets generated *in vitro* from CD34^+^ HSPC using MS5_FS12 stromal cells in the presence of human GM-CSF neutralizing antibody as compared to isotype control (n= 6 independent donors in two experiments. Line represents median).

Furthermore, we noticed that *in vitro* differentiated CD14^−^CD1c^+^ cDC2-like cells were heterogeneous for the expression of the mannose receptor CD206 (Fig. 2a). Circulating blood cDC2s do not generally express CD206 (Supplementary Fig. 3c) whereas CD206 is a marker of skin and migratory cDC2 ^19, 57^.

Most of the previously described protocols to generate human DC-like cells *in vitro* from both CD14^+^ monocytes and CD34^+^ HSPCs made an extensive use of the cytokine GM-CSF ^7, 8, 37, 39, 40, 56^, with one exception ^49^. Since we did not include GM-CSF in our protocol, we wanted to assess whether human GM-CSF was spontaneously produced in CD34^+^ cultures. We could not detect any GM-CSF from co-culture supernatant (Fig. 2b). Accordingly, GM-CSF blocking antibody did not impact the generation of cDCs driven by MS5_FS12 (Fig. 2c). We conclude that GM-CSF is dispensable for the generation of cDCs *in vitro*, as previously reported both in mouse ^58, 59^ and human ^49^.

We also observed that MS5_FS12 stromal cells significantly improve the differentiation of CD123^+^CD303/4^+^ cells (Fig. 3a), a phenotype shared by both plasmacytoid DC and pre/AS-DC ^12, 21^. A more refined phenotypic characterization of the *in vitro* generated cells also shows that all CD123^+^ cells express high levels of CD45RA, and they can be subdivided in AXL^−^ CD327^lo/-^ and AXL^+^CD327^+^ subsets, phenotypically aligning to pDC and pre/AS-DC (Fig. 3b). This conclusion was further supported by gene set enrichment analysis (GSEA) ^60^ of RNA-sequencing data, displaying a significant enrichment of previously reported pDC and AS-DC gene signatures ^21^ in *in vitro* generated AXL^−^CD327^lo/-^ and AXL^+^CD327^+^, respectively (Fig. 3c). Moreover, only the AXL^−^CD327^lo/-^ cells were capable to produce type I interferon in response to TLR stimulation, a specific feature of *bona fide* pDC which is not shared with pre/AS-DC ^12, 21^(Fig. 3d).

**Figure 3.**
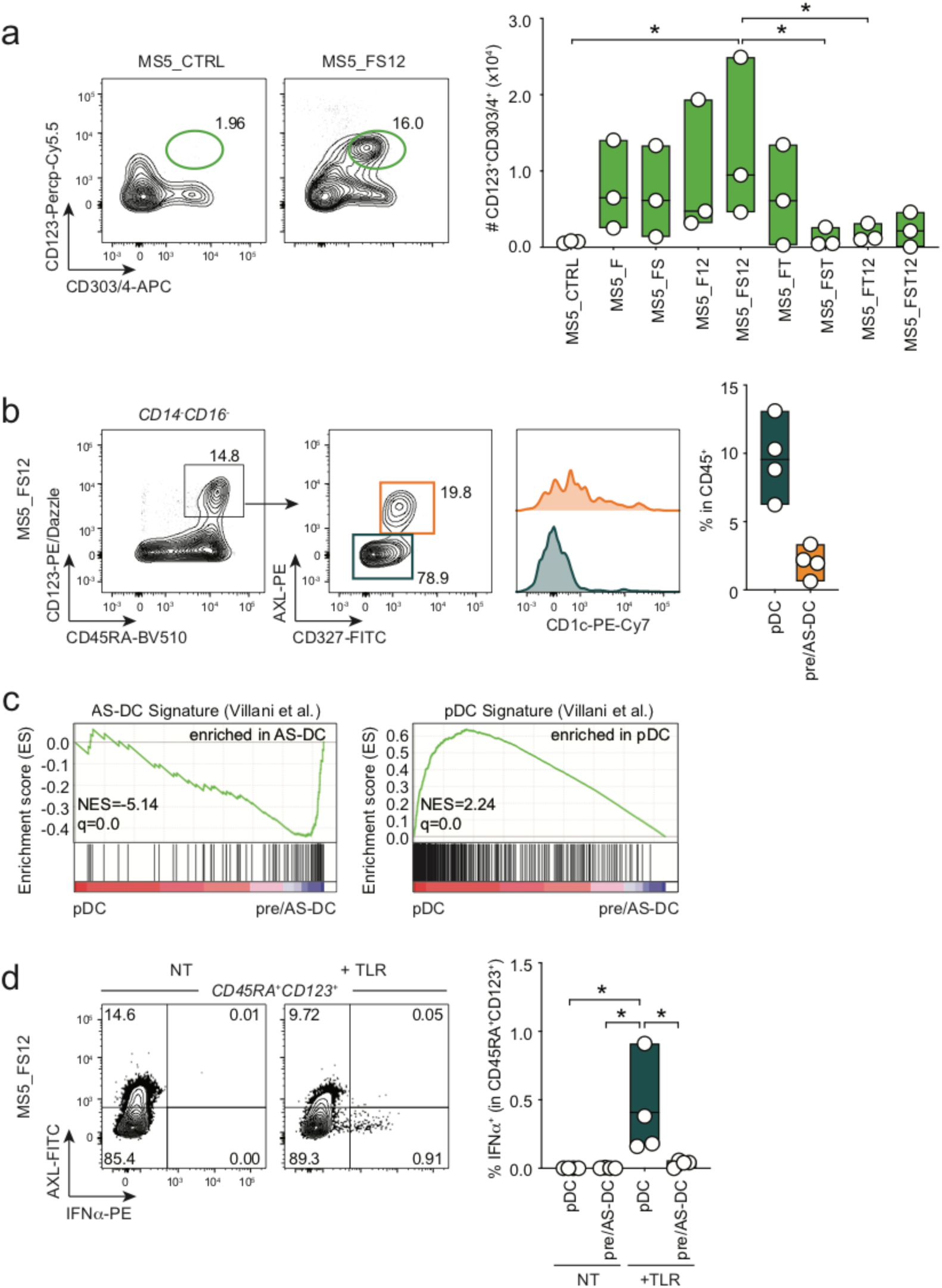
MS5_FS12 stromal cells support the development of both pDC and pre/AS-DC *in vitro*. (a) Representative FACS plots and absolute number of CD123^+^CD303/4^+^ cells generated *in vitro* from CD34^+^ HSPCs co-cultured with MS5 expressing human FLT3L (MS5_F) in combination with human SCF (S), TPO (T) and CXCL12 (12). Day15 flow cytometry analysis of n=3 cord blood donors in 3 independent experiments (line represents median; * p<0.05, one-way ANOVA test). (b) Gating strategy used to identify AXL^−^CD327^lo/-^ pDC and AXL^+^CD327^+^ pre/AS-DC within CD123^+^CD45RA^+^ cells generated *in vitro* using MS5_FS12. Graph illustrates the frequency of each subset in CD45^+^ cells (n=4 cord blood donors). (c) GSEA comparing *in vitro* differentiated pDC and pre/AS-DC using published human pDC and AS-DC gene signatures ^21^. (NES=normalized enrichment score; FDR=false detection rate). (d) Intracellular flow cytometry analysis of IFNα production in pDC and pre/AS-DC in response to 16 hours of TLR stimulation (LPS 10ng/ml, R848 1μg/ml, Poly(I:C) 25μg/ml). Bar graph shows the frequency of IFNα^+^ cells in each subset with (+ TLR) or without (NT) stimulation (n=4 cord blood donors; line represents median; * p<0.05, one-way ANOVA test).

In conclusion, we identified the combination of membrane-bound FLT3L, SCF and CXCL12 (MS5_FS12) as the most efficient tested condition to support human DCs differentiation *in vitro* from CD34^+^ HSPCs.

### Human DCs generated *in vitro* using MS5_FS12 align with circulating blood DCs

In order to validate the identity of the cDCs generated using the MS5_FS12 stromal niche, we compared the transcriptome (RNA-seq) and phenotype (CyTOF) of *in vitro* differentiated subsets to circulating blood cDC1s and cDC2s (Fig. 4a-f).

**Figure 4.**
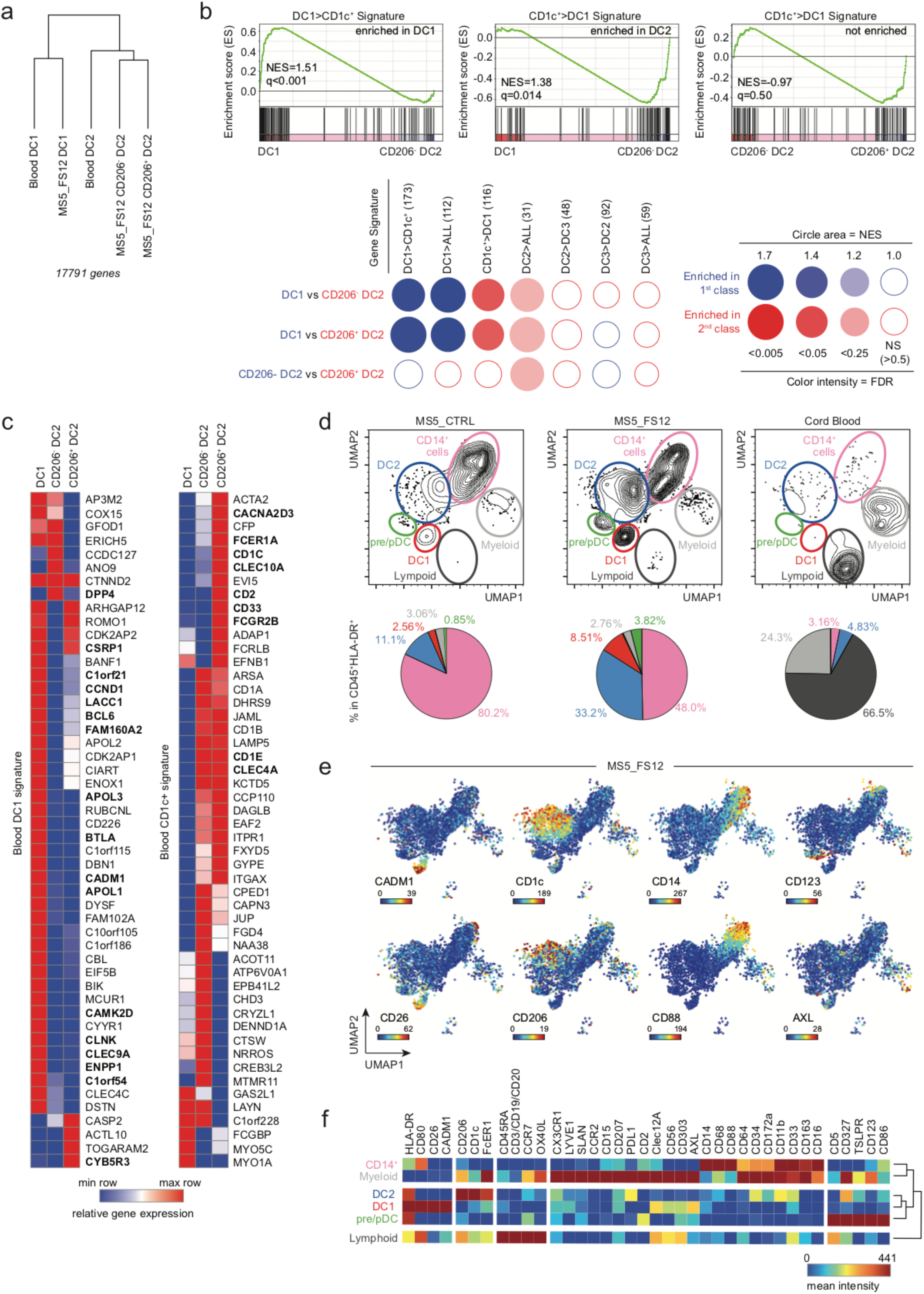
Human DCs generated *in vitro* using MS5_FS12 align with circulating blood DCs. (a) Hierarchical clustering of primary (n=3 healthy donors) versus *in vitro* generated (n=3 cord blood donors) cDCs based on 17791 genes after removing the “*in vitro* culture signature” (2000 genes) defined by pairwise comparison of primary versus *in vitro* generated subsets. (b) GSEA using blood cDC1s (DC1>CD1c^+^) and CD1c^+^ cells (CD1c^+^>DC1) signatures generated from published datasets ^63^ as well as previously published signatures of blood cDC1 (DC1>ALL), cDC2 (DC2>ALL) and cDC3 (DC3>ALL) ^21^. BubbleMap shows the enrichment of each gene signature in the pairwise comparison of CD141^+^Clec9A^+^, CD1c^+^CD206^−^ and CD1c^+^CD206^+^ cells generated *in vitro* (NES=normalized enrichment score; FDR=false detection rate). (c) Heatmaps of RNA-seq data displaying the expression of the top 50 genes of blood cDC1 and CD1c^+^ cells signatures in CD141^+^Clec9A^+^, CD1c^+^CD206^−^ and CD1c^+^CD206^+^ cells generated *in vitro*. Genes shared with previously published signatures ^21^ are highlighted in bold. (d) UMAP (Uniform Manifold Approximation and Projection) plots of CyTOF data from CD45^+^HLA-DR^+^ cells differentiated *in vitro* using MS5_FS12 and MS5_CTRL as compared to cord blood PBMCs. Pie charts indicate the frequency of each subset among the CD45^+^HLA-DR^+^ cells (mean of n=2 cord blood donors in 2 independent experiments). (e) Relative expression of selected markers in UMAP plots of CyTOF data from cells differentiated *in vitro* with MS5_FS12. (f) Heat map of markers mean intensity in each subset identified in MS5_FS12 cultures.

Hierarchical clustering of RNA-seq data revealed that subsets generated in culture maintain a strong “culture imprinting” (Supplementary Fig. 4a). Indeed, we could identify a 2000 genes signature (1000 genes up and 1000 genes down-regulated), which clearly separates *in vitro* generated cells from circulating blood subsets regardless of their subset identity (Supplementary Fig. 4b). The majority of these genes were associated to cell cycle and metabolism as shown by pathways enrichment analysis (Supplementary Fig. 4c).

Nonetheless, once this “*in vitro* culture signature” was subtracted from the total protein coding genes, CD141^+^Clec9A^+^ and CD1c^+^CD206^+/-^ cells generated in culture transcriptionally align to blood cDC1 and cDC2, respectively (Fig. 4a).

To further validate the similarity of *in vitro* generated cells with physiologically circulating subsets we performed gene set enrichment analysis (GSEA) ^60, 61^ using the BubbleGum software ^62^. This methodology enables to score the enrichment of a signature in a pairwise comparison of two transcriptomes. We scored cDC1 alignment using gene signatures specific for blood cDC1 obtained from published datasets (cDC1>CD1c^+^ ^63^ and DC1>ALL ^21^). CD14^−^ CD1c^+^ cells have recently been shown to contain two distinct subsets termed as cDC2 and cDC3 ^21^. Alignment of cultured cells was probed towards total CD1c^+^ cells (CD1c>cDC1), cDC2 (cDC2>ALL and cDC2>DC3) and DC3 signatures (DC3>cDC2 and DC3>ALL). We found that *in vitro* generated CD141^+^Clec9A^+^ and CD1c^+^CD206^+/-^ cells are enriched in genes defining circulating blood cDC1 and cDC2, respectively (Fig. 4b). The expression of the top 50 genes for each signature in the differentiated subsets further supports this conclusion (Fig. 4c). Importantly, both CD206^+^ and CD206^−^ subsets aligned preferentially to cDC2 as compared to DC3 and cDC1 (Fig. 4b and Supplementary Fig. 4d). CD163 was recently described as a marker selectively expressed in blood cDC3 as compared to cDC2 ^21^. Supporting our previous conclusion, CD163 was neither expressed in CD1c^+^CD206^−^ nor in CD1c^+^CD206^+^ cells generated *in vitro,* while CD163^+^ cells were detected among CD14^+^ monocytes and CD14^+^CD16^+^ macrophages (Supplementary Fig. 4e).

To obtain a more exhaustive characterization of the phenotype of *in vitro* generated subsets we performed CyTOF analysis using a panel of 38 metal-conjugated monoclonal antibodies. Dimension reduction of the CyTOF data was performed using the Uniform Manifold Approximation and Projection (UMAP) algorithm ^64^. UMAP plots display clusters of cells that were expanded upon MS5_FS12 co-culture as compare to MS5_CTRL (Fig. 4d). Clec9A^+^CD141^+^ cells identified by flow cytometry were shown to also express CADM1 and CD26 further aligning them with blood cDC1s (red cluster, Fig. 4d-f). CD14^−^CD1c^+^ cells did not express high level of monocyte markers such as CD64, CD68 and CD16 while they appeared heterogeneous for CD206 expression (blue cluster, Fig. 4d-f and Supplementary Fig. 4f). Of note, CD14^−^CD1c^+^ cells generated in culture did not express high level of FcεRIa, CD172a and CD5 found in blood cDC2s (Supplementary Fig. 4f). By contrast they are strongly positive for CD86 and CD80 unlike their circulating counterpart (Supplementary Fig. 4f). In addition, CD123^+^CD303^+^ cells were shown to express heterogeneous levels of pre-DC markers such as CD327 and CX3CR1 and moderate level of AXL (green cluster, Fig. 4d-f and Supplementary Fig. 4f), in line with flow cytometry analysis highlighting the presence of both pDC and pre/AS-DC within CD123^+^CD303^+^ cells generated *in vitro* (Fig. 3b). On the other hand, by combining flow and mass cytometry analysis we were able to show that MS5_FS12 stromal cells do not support lymphoid development (Fig. 4d and Supplementary Fig. 4g). Indeed, the remaining cells (other than DC) present in culture consist of CD15^+^ Granulocytes and CD14^+^CD16^+/-^ Monocytes/Macrophages (Supplementary Fig. 4h). Finally, the analysis of *in vitro* cDC differentiation kinetics revealed that both cDC1 and cDC2 can be detected in MS5_FS12 cultures as early as day7 (Supplementary Fig. 4i). However, the yield of *in vitro* generated cDC was significantly higher at day14, when most of the cultures were therefore analyzed (Supplementary Fig. 4i).

Collectively, our data demonstrate that: *i) in vitro* generated CD141^+^Clec9A^+^ recapitulate the phenotype of *bona fide* blood cDC1; *ii)* CD14^−^CD1c^+^ cells align to cDC2 regardless of their CD206 expression; *iii)* CD123^+^CD303^+^ cells contain some recently described pre-DC/AS-DC phenotypically and functionally distinct from pDCs. However, we identified two major limitations of the *in vitro* culture. First, the culture system imposes a strong transcriptional imprinting throughout subsets. Second, *in vitro* generated cDC2s failed to express to full phenotypic profile of blood cDC2s.

### Subcutaneous engraftment of MS5_FS12 in NSG mice results in the formation of a “synthetic niche” supporting human CD34^+^ progenitor local maintenance, differentiation and expansion

We next wanted to assess whether we could use MS5_FS12 to recapitulate a more physiological niche microenvironment supporting human HSPCs maintenance *in vivo*.

To this end, we designed an experimental strategy based on the subcutaneous injection of cord blood-derived CD34^+^ HSPCs together with MS5_FS12 in a basement membrane matrix (Matrigel) in NSG mice (Fig. 5a).

**Figure 5.**
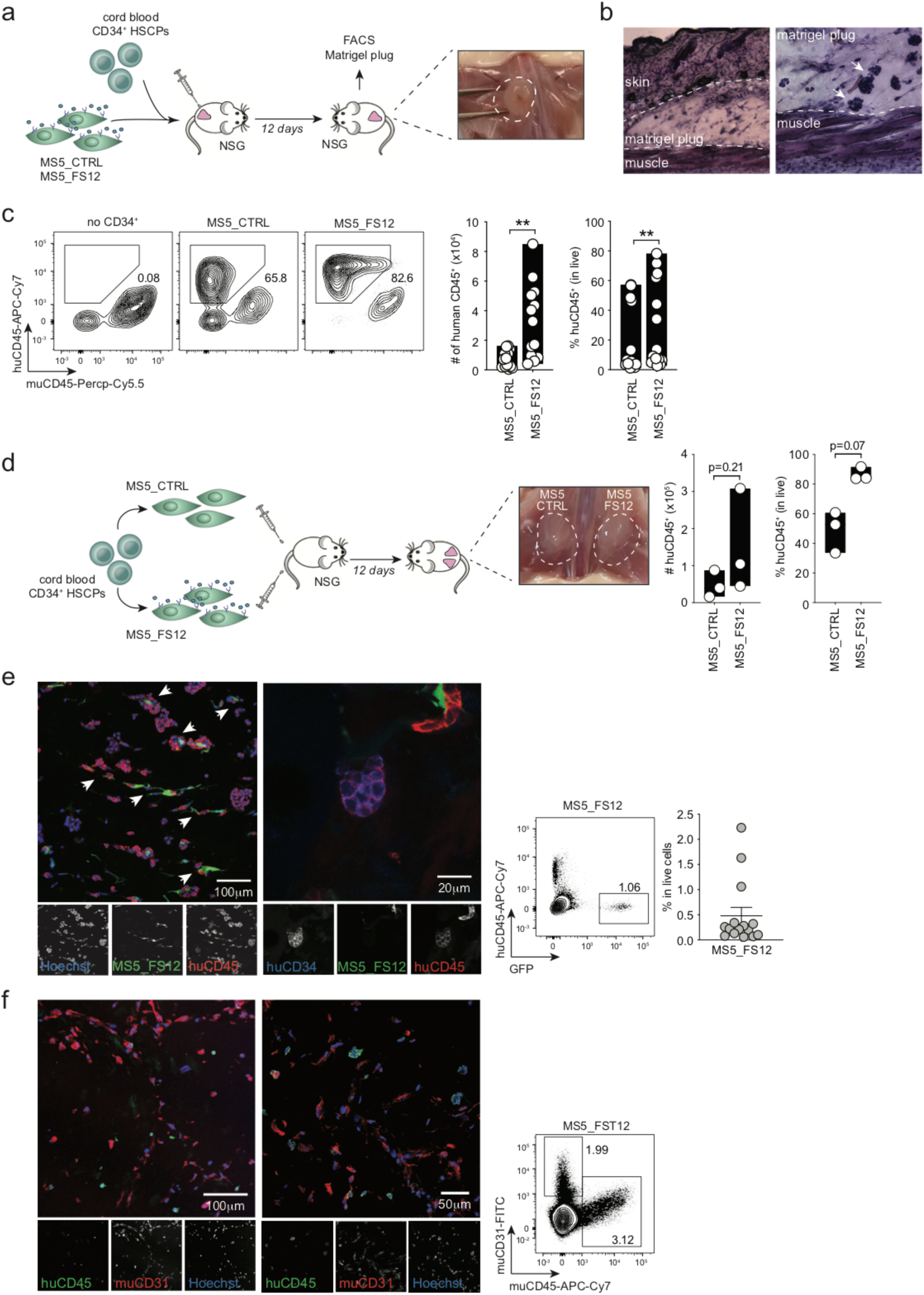
Subcutaneous engraftment of MS5_FS12 in NSG mice results in the formation of a “synthetic niche” supporting human CD34^+^ progenitor local maintenance, differentiation and expansion. (a) Experimental strategy for an *in vivo* synthetic niche. Human HSPCs were injected subcutaneously along with MS5_FS12 in a basement membrane matrix (Matrigel) preparation. (b) Hematoxylin-eosin staining of subcutaneous organoids at day 12. Arrows show clusters of Matrigel-embedded cells. (c) Flow cytometry analysis at day 12 of Matrigel organoids containing either MS5_CTRL or MS5_FS12 cells. Absolute number and frequency of human CD45^+^ cells recovered are summarized in bar graphs (n=13 cord blood donors in 6 independent experiments; ** p<0.01, two-tailed paired Student t test). (d) Experimental strategy and quantification of human CD45^+^ cells recovered from physically separated plugs containing either MS5_CTRL or MS5_FS12 cells injected in the same recipient (n=3 cord blood donors in one experiment; two-tailed paired Student t test). (e) Immunofluorescence staining of plug sections displaying the interaction of GFP^+^ MS5_FS12 (green) with human CD45^+^ cells (red). Human hematopoietic progenitors were also identified as CD45^+^ (red) CD34^+^ (blue) cells in MS5_FS12 plugs. Nuclei were stained with Hoechst (blue). Arrows show interaction of human CD45^+^ leukocytes with GFP^+^ MS5_FS12. The presence of GFP^+^ stromal cells in Matrigel organoids at day 12 was further confirmed by flow cytometry. (f) Visualization of mouse CD31^+^ endothelial cells by immunofluorescence. Fixed sections were stained for human CD45 (green) and mouse CD31 (red). Nuclei were stained with Hoechst (blue). The presence of mouse CD31^+^ cells was further confirmed by flow cytomery.

Clusters of cells embedded in Matrigel can be identified as early as day 12 by tissue histology (Fig. 5b). Flow cytometry analysis demonstrated that MS5_FS12 but not MS5_CTRL induced the expansion of human leukocytes within the Matrigel plugs (Fig. 5c). We then tested whether cell-to-cell interactions of eMSC with human progenitors play a role in this process. We injected two independent plugs of CD34^+^ HSPCs with either MS5_CTRL or MS5_FS12 in the same recipient mouse (contralateral plugs) (Fig. 5d). We found a relative expansion of human leukocytes in MS5_FS12 as compared to MS5_CTRL contralateral plugs (Fig. 5d). We conclude that MS5_FS12 does not efficiently provide soluble factors enabling human leukocytes expansion systemically. Therefore, we hypothesized that membrane bound FLT3L and SCF together with CXCL12 define an efficient *in vivo* niche by delivering cell-to-cell contacts supporting HSPCs expansion. Supporting this hypothesis, we found that MS5_FS12 expressing GFP persist in the plugs at day 12 of differentiation (Fig. 5e). Immunofluorescence analysis further supported this observation and demonstrated the existence of cell-to-cell contact between MS5_FS12 and human CD45^+^ leukocytes (Fig. 5e and Supplementary Fig. 5a). Leukocytes expressing CD34^+^ could also be detected, supporting the notion that a pool of undifferentiated progenitors is maintained in the MS5_FS12 organoids at day 12 (Fig. 5e). Of note, Matrigel plugs contained some mouse CD31^+^ cells suggesting undergoing vascularization as evidenced by the formation of early tube-like structure (Fig. 5f). However, no vascular leak was observed, as demonstrated by the absence of intravenously delivered CTV^+^CD3^+^ cells in the subcutaneous plug (Supplementary Fig. 5b).

Taken together, these data show that engineered stromal cells MS5_FS12 provide a minimal synthetic niche scaffold supporting human CD34^+^ HSPCs maintenance and expansion *in vivo*.

### The MS5_FS12 niche efficiently supports human cDC1, cDC2, pre/AS-DC and pDC development from CD34^+^ HSPCs *in vivo*

We investigated whether the engraftment of CD34^+^ HSPCs together with MS5_FS12 could lead to the local differentiation of human DC subsets.

Flow cytometry analysis of Matrigel organoids demonstrates that the MS5_FS12 but not the MS5_CTRL niche specifically supports the differentiation of CD141^+^Clec9A^+^ cDC1-like cells and CD14^−^CD1c^+^ cDC2-like cells (Fig. 6a and Supplementary Fig. 6a). This finding was supported by immunofluorescence staining on plug sections highlighting the occurrence of human CD45 cells expressing either Clec9A or CD1c (Fig. 6b).

**Figure 6.**
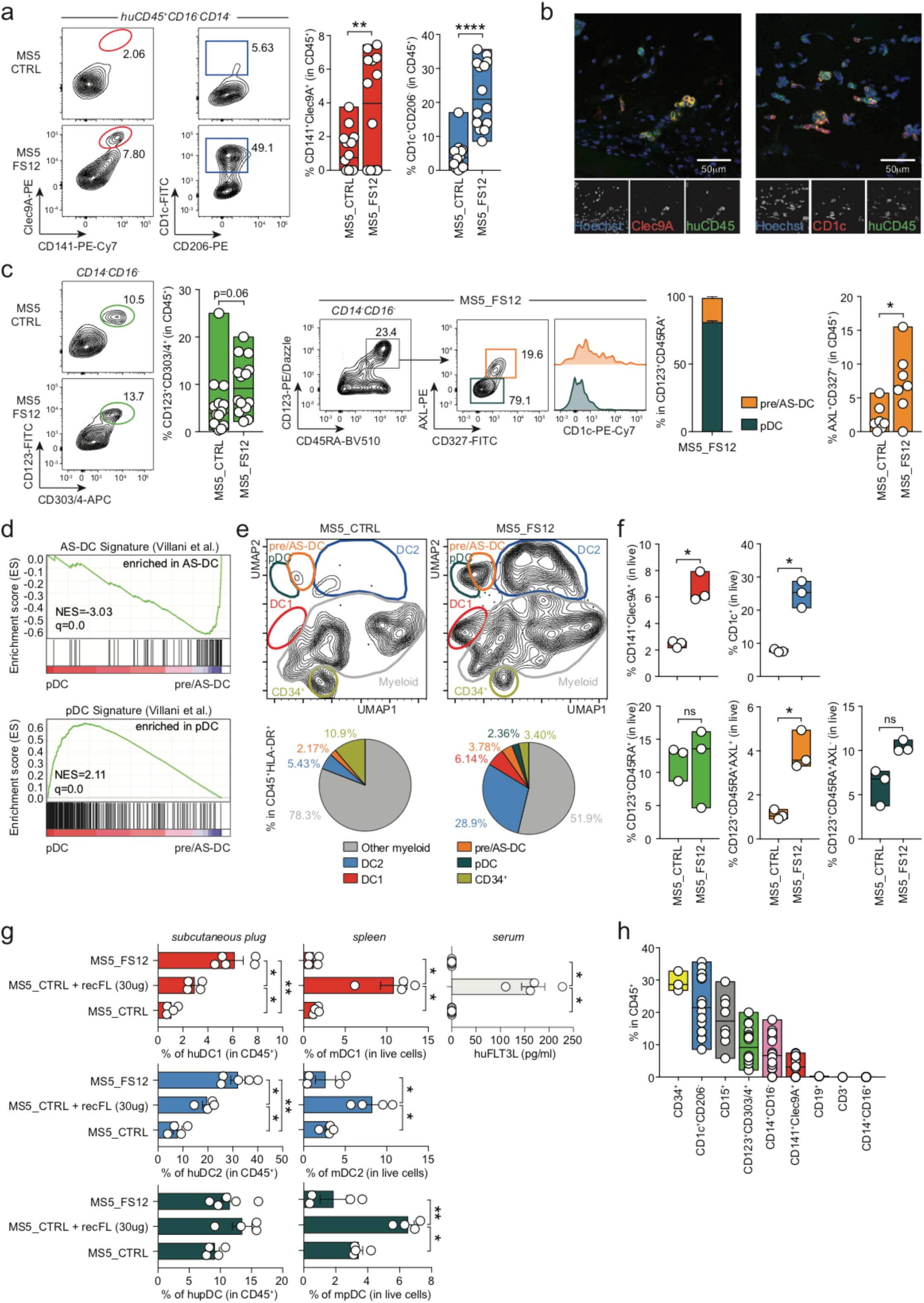
The MS5_FS12 niche efficiently supports human cDC1, cDC2 and pre/AS-DC development from CD34^+^ HSPCs *in vivo*. (a) Flow cytometry analysis of Matrigel organoids containing either MS5_CTRL or MS5_FS12 stromal cells. Bar graphs show the frequency of CD141^+^Clec9A^+^ cDC1 and CD1c^+^CD206^−^ cDC2 in total CD45^+^ cells (n=14 donors in 6 independent experiments. Line represents median; **p<0.01 ****p<0.0001, two-tailed paired Student t test). (b) Immunofluorescence staining of Matrigel plugs sections confirming the presence of huCD45^+^ (green) Clec9A^+^ (red) cDC1 and huCD45^+^ (green) CD1c^+^ (red) cDC2 in MS5_FS12 organoids *in vivo*. Nuclei (blue) were stained with Hoechst. (c) *(left panel)* Flow cytometry analysis and quantification of CD123^+^CD303/4^+^ cells recovered from MS5_CTRL and MS5_FS12 organoids (n=14 donors in 6 independent experiments. Line represents median; two-tailed paired Student t test). *(middle panel)* Gating strategy used to identify AXL^−^CD327^lo/-^ pDC and AXL^+^CD327^+^ pre/AS-DC within CD123^+^CD45RA^+^ cells generated *in vivo* in MS5_FS12 organoids. Bar graph illustrates the frequency of each subset in CD123^+^CD45RA^+^ cells (n=3 cord blood donors). *(right panel)* Frequency of pre/AS-DC in total CD45^+^ cells in MS5_CTRL vs MS5_FS12 organoids (n=7 donors in 4 independent experiments. Line represents median; * p<0.05, two-tailed paired Student t test). (d) GSEA comparing *in vivo* differentiated pDC and pre/AS-DC using published human pDC and AS-DC gene signatures ^21^. (NES=normalized enrichment score; FDR=false detection rate). (e) UMAP plots of CyTOF data comparing CD45^+^HLA-DR^+^ cells differentiated *in vivo* using MS5_FS12 and MS5_CTRL. Pie charts indicate the frequency of each subset among the CD45^+^HLA-DR^+^ cells (mean of n=2 cord blood donors in 2 independent experiments). (f) Frequency of CD141^+^Clec9A^+^ cDC1, CD1c^+^CD206^−^ cDC2, CD123+CD45RA+AXL-pDC, CD123+CD45RA+AXL+ pre/AS-DC and total CD123^+^CD45RA^+^ cells recovered from two physically separated plugs containing either MS5_CTRL or MS5_FS12 injected in the same recipient (n=3 cord blood donors in one experiment; line represents median; * p<0.05, two-tailed paired Student t test). (g) To assess the local versus systemic effect of MS5_FS12 niche, NSG mice were injected subcutaneously either with MS5_CTRL or MS5_FS12 stromal cells. Human recombinant FLT3L was administered intra-peritoneum to mice bearing MS5_CTRL plugs at day 0, 4 and 8 (10ug/mouse/injection) (MS5_CTRL+recFL). Frequency of *in vivo* differentiated human cDC1, cDC2 and pDC detected in subcutaneous organoid (left) and murine cDC1, cDC2 and pDC detected in the spleen (center) were reported. The levels of human recombinant FLT3L in the serum were measured by ELISA (right). Mean ± SEM of n=4 mice/group in 2 independent experiments. * p<0.05, **p<0.01, one-way ANOVA test. (h) Frequency of *in vivo* differentiated cell subsets among the total huCD45^+^ cells recovered from MS5_FS12 plugs at day 12 (line represents median).

Further analysis revealed the expansion of CD123^+^CD303/4^+^ cells in MS5_FS12 when compared to MS5_CTRL plugs (Fig. 6c and Supplementary Fig. 6a). All these cells also expressed CD45RA and heterogeneous levels of AXL and CD327, as previously described for their *in vitro* counterparts (Fig. 6c). However, only MS5_FS12 induced a strong accumulation of AXL^+^CD327^+^ pre/AS-DC expressing various levels of CD1c (Fig. 6c). In addition, *bona fide* CD123^+^CD45RA^+^AXL^−^CD327^lo/-^ pDCs could also be detected (Fig. 6c). RNA-seq analysis of *in vivo* generated CD123^+^AXL^−^CD327^lo/-^ and CD123^+^AXL^+^CD327^+^ cells further support this conclusion and unequivocally align them to blood circulating pDC and AS-DC, respectively (Fig. 6d).

To further refine the phenotypic characterization of HLA-DR^+^ mononuclear phagocytes in MS5_FS12 organoids we performed high-dimensional mass cytometry analysis. The comparison of MS5_FS12 with MS5_CTRL plugs highlighted the expansion of all subsets previously identified by flow cytometry: cDC1s, cDC2s, pre/AS-DCs, pDCs and a distinct population of CD33^+^CCR2^+^CX3CR1^+^Clec12A^+^ myeloid cells (Fig. 6e and Supplementary Fig. 6b). We next wanted to determine whether commitment towards the cDC lineage would be dependent on local developmental cues and possibly cell-to-cell contact between CD34^+^ HSPCs and MS5_FS12. To this end, we engrafted mice with two distal organoids, one containing MS5_CTRL and the second one containing MS5_FS12. We found that cDC1, cDC2 and pre/AS-DCs were selectively expanded in MS5_FS12 plugs (Fig. 6f and Supplementary Fig. 6c). On the contrary, pDC were not significantly increased in the same comparison (Fig. 6f and Supplementary Fig. 6c). We conclude that local cues associated to the MS5_FS12 niche control cDC lineage commitment. In support of this view, we could not detect a systemic increase in the levels of serum FLT3L in mice carrying engineered stromal cell plugs (Fig. 6g). Accordingly, spleen resident murine cDCs did not expand upon MS5_FS12 engraftment while they massively do so upon administration of recombinant soluble human FLT3L (Fig. 6g). Together with the 2-plugs experiments (Fig. 6f), these observations suggest that most of the FLT3L aegis relies on its membrane bound form delivered in the context of eMSCs. Of note, administration of recombinant soluble FLT3L was poorly efficient at expanding human DCs populations in Matrigel organoids formed with control stromal cells (Fig. 6g). This demonstrates the superiority of local, cell-associated cues (MS5_FS12, *i.e*.) to achieve the expansion of human cDCs in the dermis of NSG mice.

A more extensive characterization of MS5_FS12 organoids revealed the presence of myeloid lineages other than DCs, such as CD14^+^CD16^−^ monocyte-like cells and CD15^+^ granulocytes (Fig. 6h and Supplementary Fig. 6d). Conversely, no lymphoid specification was observed (Fig. 6h and Supplementary Fig. 6d). Despite this broad spectrum of lineages, CD34^+^ HSPCs represented the most abundant population at day 12 (Fig. 6h). This observation suggests that the MS5_FS12 niche combines HSPC maintenance with lineage commitment.

### cDC2 generated *in vivo* more faithfully align to blood circulating cDC2s

Finally, we wanted to establish whether *in vivo* differentiated DCs in MS5_FS12 organoids had a distinct phenotype from the subsets generated *in vitro* in MS5_FS12 co-culture.

UMAP plots of CyTOF analysis revealed three major findings. First, pre/AS-DCs represent a more abundant population *in vivo* (Fig. 7a and Supplementary Fig. 7a). Second, both cDC1 generated *in vitro* and *in vivo* fully align phenotypically, displaying a strong expression of CADM1 and CD26 (Fig. 7a and Supplementary Fig. 7a). Third, unlike cDC1, cDC2 generated *in vivo* exhibit noticeable phenotypic differences. *In vivo*-generated cDC2 express higher levels of FcεRIa, CD172a and CD5 while showing lower expression of HLA-DR and CD86 (Fig. 7b and Supplementary Fig. 7b). The specific phenotype conferred by the MS5_FS12 niche education renders cDC2s more akin to their blood counterparts. In order to compare extensively the transcriptional landscape of *in vivo* (NSG organoids) generated DCs with primary DCs found in human blood, we performed RNAseq analysis on FACS-sorted cDC2s obtained after the enzymatic digestion of MS5_FS12-containing plugs or purified from human blood.

**Figure 7.**
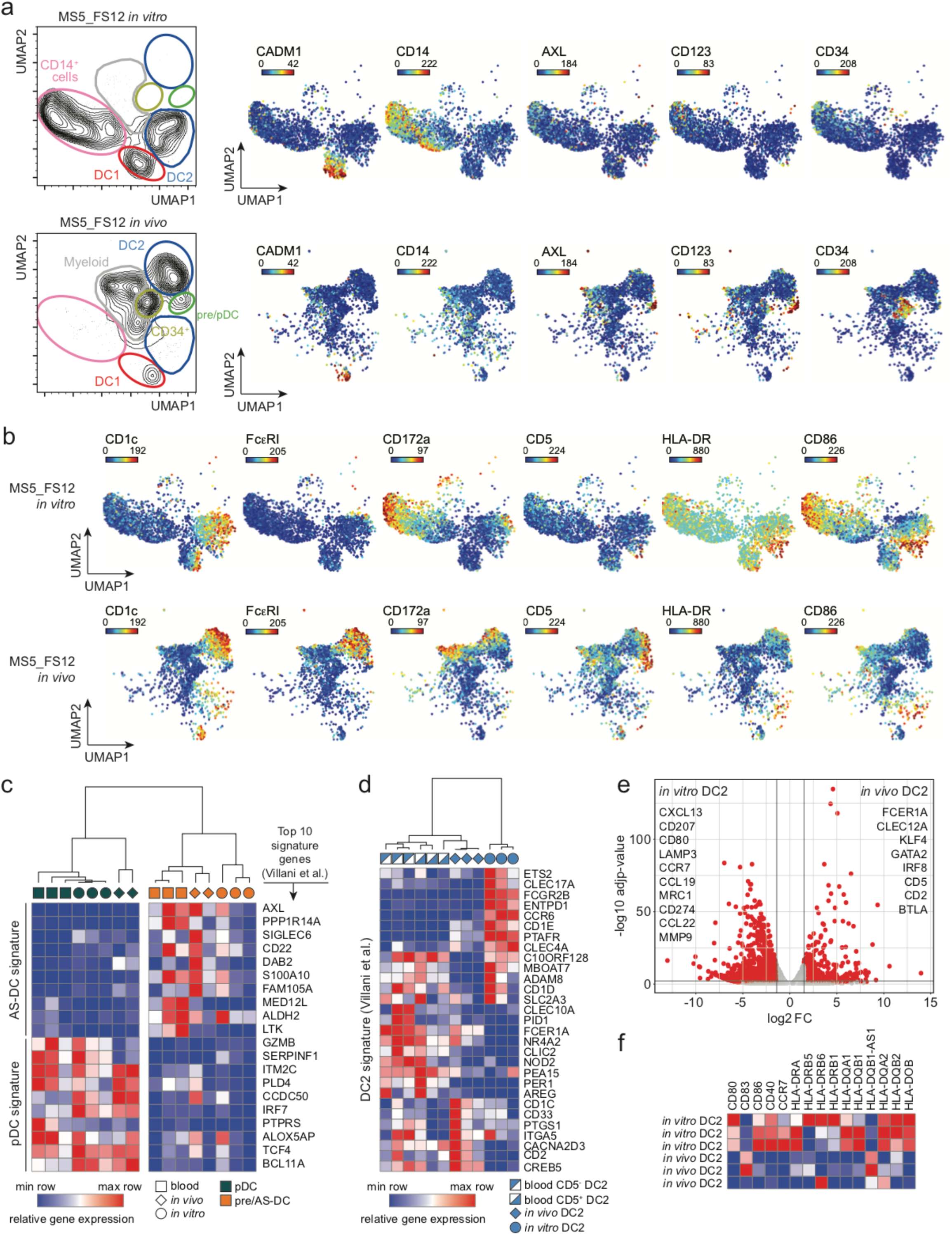
*In vitro* generated cDC2s more faithfully recapitulate the phenotype of human circulating cDC2s. (a) UMAP plots of CyTOF data comparing CD45^+^HLA-DR^+^ cells generated using MS5_FS12 stromal cells either *in vitro* or *in vivo*. Relative expression of selected markers is shown for each condition. (b) Relative expression of selected markers highlighting the phenotypic differences between cDC2s generated *in vitro* and *in vivo* using MS5_FS12 stromal cells. (c) Heatmap displaying gene expression of the top 10 genes of blood pDC and AS-DC published signatures ^21^ in pDC and pre/AS-DC generated *in vitro*, *in vivo* and isolated from blood PBMC (n=2-3 independent donors). (d) Heatmap displaying gene expression of the blood cDC2 published signature ^21^ in cDC2 cells generated *in vitro*, *in vivo* and isolated from blood PBMC (n=3 independent donors). (e) Volcano plot showing differentially expressed genes between *in vitro* and in vivo generated cDC2 (Log2FC>1.5, adjusted p-value<0.05). (f) Heatmap displaying gene expression of activation markers and co-stimulatory molecules expressed in cDC2 generated *in vivo* and *in vitro* (n=3 independent donors).

As previously observed for *in vitro* generated cells, the MS5_FS12 niche confers an *in vivo* imprinting resulting in the differential expression of 2872 genes (up- or down-regulated) in *in vivo* versus *ex vivo* isolated subsets (Supplementary Fig. 7c). Pathway analysis revealed that this *in vivo* bias was mainly due to upregulation of genes associated with DNA replication, cell cycle and proliferation (MYC, CDC6/7, POLA2, MCM6/7 e.g.) (Supplementary Fig. 7c and Supplementary Table 6).

Moreover, we found that: *i*) AXL^+^Siglec6^+^ pre/AS-DCs generated *in vivo* (or *in vitro*) align to their primary counterparts and selectively express a signature that distinguish them from *bona fide* pDCs (*DAB2, CD22, ADLH2* e.g.) (Fig. 7c). *ii*) conversely, AXL^−^Siglec6^−^ *bona fide* pDCs generated *in vivo* (or *in vitro*) align to their primary counterparts and express high levels of markers distinguishing them from pre/AS-DCs (*IRF7, GZMB, TCF4, BCL11A, e.g.*) (Fig. 7c). *iii*) cDC2s generated *in vivo* (in NSG mice organoids carrying MS5_FS12) had higher levels of similarity with blood cDC2s (including higher expression of *BTLA*, *FCER1A,* e.g.) (Fig. 7b, 7d and 7e). Recently, both CD5^+^ and CD5^−^ cDC2s subsets have been reported in human blood ^65, 66^ and we found that *in vivo* generated cDC2s aligned particularly well with blood CD5^+^ cDC2s (with the expression of CD5, CD2, e.g.) (Fig. 7b and 7e). By contrast, *in vitro* generated cDC2s expressed high levels of activation genes such MHC molecules (*HLA-DR, DQ*, *e.g*.); co-stimulatory molecules (*CD80, CD40*, *e.g*.), activation markers (*ETS2, CCR6, CCR7, CXCL13, CCL22*, *e.g*.) (Fig. 7e and 7f) and genes associated with type I and type II interferon pathways (*STAT1, IRF9, IGS15, GBP1,* e.g.) (Supplementary Fig. 7d and Supplementary Table 6).

All together, we conclude that MS5_FS12-containing organoids provide a unique scaffold for the specification and commitment of the DC lineage. This unique and versatile system bypasses the limitation of *in vitro* cultures, which generated inefficiently pre/AS-DCs and biased the differentiation of cDC2s toward an activated phenotype. Collectively, MS5_FS12 organoids faithfully recapitulate the differentiation of DCs with an unattained level of similarity to human blood DCs, especially blood cDC2s.

### cDC2 generated *in vitro* and *in vivo* recapitulate functional features ascribed to blood cDC2

In the last set of experiments, we aimed at functionally validate cord blood-derived cDC generated in the MS5_FS12 stromal niche. Moreover, we also assessed whether the phenotypic differences observed in cDC2 generated *in vitro* and *in vivo* may impact their function.

We first confirmed *in vitro* the responsiveness of cord blood-derived cDC2 to TLR agonists expressed in human circulating cDC2, as demonstrated by the up-regulation of maturation markers (i.e. HLA-DR, CD86 and CD83) in response to TLR4 (LPS) and TLR8 (VTX) stimulation (Fig. 8a). We then performed a mixed leukocyte reaction (MLR) by co-culturing CTV-labeled allogeneic naïve T cells together with FACS-sorted cDC subsets (Supplementary Fig. 8a) activated overnight with a TLR agonists cocktail comprising of LPS (TLR4), R848 (TLR7/8) and Poly(I:C) (TLR3). After 7 days of culture, we observed that both *in vitro* and *in vivo* generated cDC2 and pre/AS-DC were capable to efficiently induce CD4^+^ naïve T cells proliferation (Fig. 8b), as expected and reported for circulating blood cDC2 ^12, 21^ (Supplementary Fig. 8b). Conversely, pDC were significantly less effective on triggering T cells activation, as shown by the consistent reduction in the frequency of dividing CD4^+^ T cells when compared to cDC2 and pre/AS-DC (Fig. 8b).

**Figure 8.**
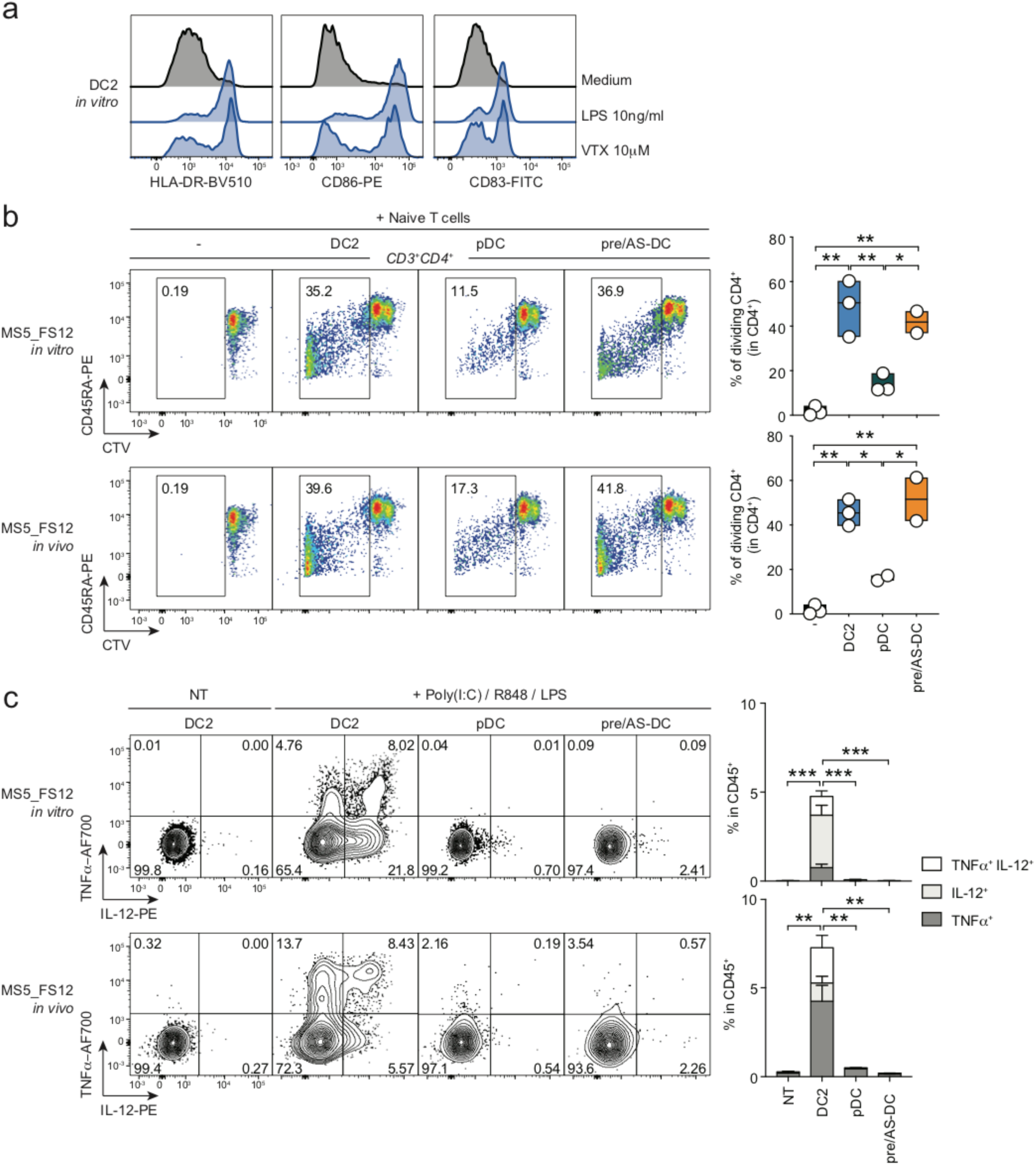
cDC2 generated *in vitro* and *in vivo* recapitulate functional features ascribed to blood cDC2. (a) Histograms showing up-regulation of activation markers HLA-DR, CD86 and CD83 in *in vitro*-differentiated cDC2 in response to TLR4 (LPS) and TLR8 (VTX) overnight (16 hours) stimulation. (b) Representative FACS plots and quantification of mixed lymphocyte reaction (MLR) using *in vitro* and *in vivo* differentiated cDC2, pDC and pre/AS-DC. FACS-sorted DC subsets were activated overnight (16 hours) using a TLR agonist cocktail (LPS 10ng/ml, R848 1μg/ml and Poly(I:C) 25μg/ml) and co-cultured with CTV-labeled naïve T cells for 7 days; n=2-4 independent cord blood donors in two independent experiments; line represents median; * p<0.05, **p<0.01, one-way ANOVA test. (c) Intracellular flow cytometry analysis of TNFα and IL-12 expression in *in vitro* and *in vivo*-differentiated DC subsets upon overnight (16 hours) stimulation with TLR agonist cocktail (LPS 10ng/ml, R848 1μg/ml and Poly(I:C) 25μg/ml). Mean ± SEM of n=4 independent cord blood donors. ** p<0.01, ***p<0.001, one-way ANOVA test.

Importantly, only cDC2 were able to produce high amounts of T cell-polarizing cytokines in response to TLR stimulation, as demonstrated by intracellular TNFα and IL-12 detection by flow cytometry (Fig. 8c). All these features demonstrate that cDC2s, pDCs and pre/AS-DCs functionally align to their *in vivo* counterparts as previously described in the literature ^12, 21^.

Collectively, our data suggest that: *i*) both *in vitro* and *in vivo* differentiated cDC2 are equally capable to induce CD4^+^ T cells activation as well as produce large amounts of TNF-α and IL-12; *ii*) pre/AS-DC are as efficient as cDC2 in activating allogeneic naïve CD4+ T cells *in vitro*, a distinctive feature that clearly separate them from the pDC lineage; *iii*) despite their ability to induce CD4^+^ T cell proliferation in MLR settings, pre/AS-DC do not produce high levels of cytokines commonly associated with cDC2 function, such as TNF-α or IL-12.

## Discussion

Over the last two decades, DC-based strategies have been proposed for the therapeutic vaccination against cancer, including (*i*) non-targeted protein-based vaccines captured by DCs *in vivo*, (*ii*) specific targeting of DC subsets with mAb coupled to tumor antigens ^67^ and (*iii*) antigen loading of *ex vivo* generated dendritic cells ^3^. In this context, experimental models recapitulating the development of human DC subsets are crucially needed.

Here we describe a novel approach to model human DC development from CD34^+^ HSPCs both *in vitro* and *in vivo*. To this end, we primarily focused on the physiological niches where human DCs differentiate and maintain: a central bone marrow niche where DC progenitors are specified and peripheral niches in the lymph nodes where DCs reside.

Previous studies have shown that the cell-to-cell interaction with membrane-bound factors expressed by the niche microenvironment plays an essential role in HSPCs maintenance and expansion ^22, 23, 68, 69^. Alternative splicing of human and murine SCF transcript results in the synthesis of both a soluble and a membrane-bound non-cleavable form of the protein. Interestingly, the secreted form of SCF/KITLG is not sufficient for the establishment of a functional niche in murine bone marrow ^22, 23^ whereas the expression of human membrane-bound SCF is sufficient to support human myeloid development in humanized mice ^69^. We therefore wanted to test whether a similar relationship might exist between the soluble and membrane-bound forms of human FLT3L. Consistent with this hypothesis, the expression of transmembrane FLT3L in mesenchymal stromal cells (MS5_F) improved the efficiency of DC differentiation *in vitro*, as compared to its soluble form (MS5+recFL). Moreover, the engraftment of distal organoids (MS5_CTRL vs MS5_FS12) together with the comparison of local (membrane bound) vs systemic (soluble) delivery of human FLT3L *in vivo*, supported the notion that cell-associated FLT3L delivered by engineered stromal cells significantly improves the development of human DCs in NSG mice.

Several protocols have been proposed for the *in vitro* differentiation of human cDCs from CD34^+^ HSPCs ^7, 38–40, 48, 49^. *In vitro* differentiated cDC1s have been shown to fully recapitulate both the phenotype and function of circulating *bona fide* blood cDC1s ^8, 39, 48, 49^ including high levels of IRF8 expression and Batf3-dependency *in vitro* ^13^, as well as IRF8-dependancy both *in vivo* ^70^ and *in vit*ro ^48^. Conversely, several aspects have limited an exhaustive validation of *in vitro* generated CD1c^+^ cDC2-like cells such as the expression of CD14 ^7^, the transcriptional alignment with monocyte-derived DCs (moDCs) ^39^ or the lack of a high-dimensional phenotypic characterization ^48, 49^.

Here we described a new system (MS5_FS12), which efficiently supports the differentiation of both CD141^+^Clec9A^+^ cDC1s and CD14^−^CD1c^+^ cDC2s. However, cDC2s generated in MS5_FS12 cultures only partially recapitulate the phenotype of circulating blood cDC2s, as suggested by the lack of expression of cDC2-specific markers such as FcεRIa and CD5. Nevertheless, engineered MSCs display the unique advantage of being suitable for *in vivo* applications.

Immunodeficient mice provide a unique system to model the onset of human immune responses in realistic settings ^71^. However, a reliable method to achieve the differentiation of human DCs *in vivo* has not been described, yet. Current protocols rely on the engraftment of human CD34^+^ HSPCs in sub-lethally irradiated immunodeficient mice (humanized mice). This strategy has not been successful in the generation of well-characterized circulating DC subsets ^71^. Administration of supraphysiological levels of recombinant FLT3L has been shown to stimulate cDC differentiation upon reconstitution of NSG ^29, 50^ or *Flt3*^−/−^ BRGS ^51^ mice with human CD34^+^ HSPCs. However, the phenotype of CD141^+^ cDC1s and CD1c^+^ cDC2s was poorly characterized and, despite exceptions ^52^, tissue DCs were not generally investigated. These aspects represent an important limitation by precluding, for instance, the modeling of skin vaccination. Alternatively, transgenic mice expressing human GM-CSF and IL-3 (in the presence or absence of human SCF), either constitutively ^72, 73^ or by replacing their murine counterparts (knock-in) ^74^, have been generated. Despite displaying higher levels of myeloid reconstitution, as well as the presence of human alveolar macrophages in the lungs of humanized mice ^74^, this approach did not improve the development of human cDCs in lymphoid and non-lymphoid tissues of engrafted animals.

We demonstrated that MS5_FS12 support the differentiation of human cDCs *in vivo* in subcutaneous organoids in NSG mice. High-dimensional mass cytometry (Cytof) and transcriptomic (RNA-seq) analysis of *in vivo* generated cells confirmed their phenotypic and transcriptional alignment to circulating blood cDC1s and cDC2s. More importantly, cDC2s generated *in vivo* better resemble their physiologically circulating counterparts by expressing higher levels of FcεRIa, CD172a, CD5, CD2 and BTLA when compared to *in vitro* differentiated cells. The lower expression of activation markers such as CD86, CD80 and MHC molecules also suggests that *in vivo* cDC2s displayed a less mature phenotype than *in vitro* generated cells.

Moreover, MS5_FS12 niche was capable of supporting the local maintenance and expansion of human HSPCs as well as pre/AS-DCs, resulting in the persistence of a long-lasting source of progenitors capable of undergoing DC differentiation. To our knowledge, this is the first time that a well-characterized system supporting the development of human pre/AS-DC is reported.

Collectively, we have demonstrated that the engineered stromal cells MS5_FS12 give rise to a synthetic hematopoietic niche when injected subcutaneously in NSG mice. The niche microenvironment efficiently supports the expansion of CD34^+^ HSPCs, and human DCs subsets (cDC1, cDC2 and pre/AS-DC) can be detected as early as day 12 in a radiation-free environment. Importantly, *in vitro* culture system imposes a certain level of spontaneous activation that is not found in primary circulating blood DCs. Differentiation of human cDCs within humanized mice limit this phenomenon and make cDCs more similar to circulating primary cDCs. Hence, this approach represents a versatile system to study human DC development and function *in vivo*.

## Methods

### Mice

All *in vivo* experiments were performed using NOD.Cg-*Prdc^scid^ Il2rg^tm1Wjl^*/SzJ (NSG) mice (JAX #005557). All mice were used between 8 and 12 weeks of age. They were maintained in specific-pathogen-free conditions and handled according to protocols approved by the UK Home Office.

### Generation of engineered MSCs

Human FLT3L, SCF and TPO were amplified by PCR from cDNA expression plasmids (Origene) and cloned into pMX retroviral vectors (vectors details in Supplementary Table 1). Lentiviral vector pBABE-puro-SDF-1 alpha was a gift from Bob Weinberg (Addgene plasmid #12270) ^75^. Viral particles were generated using the retroviral packaging plasmid pCL-Ampho and a second generation lentiviral packaging system (psPAX2 and pMD2.G), respectively. MS5 cells were first transduced with CXCL12 lentiviral vector and selected using 15 μg/ml of Puromycin (Thermo Fisher). Then, a combination of single or multiple cytokines were used to transduce MS5 cells as illustrated in Supplementary Fig. 2a. Cells expressing human membrane bound FLT3L and SCF were sorted according to antibody staining of the transmembrane proteins (antibodies listed in Supplementary Table 2). TPO-expressing cells were sorted according to the expression of mCherry reporter.

### Flow cytometry and fluorescent-activated cell sorting (FACS)

Extracellular staining of cells was preformed in FACS buffer, consisting in PBS (Life Technologies), 1% BSA (Apollo Scientific) and 2 mM EDTA (Life Technologies). For intracellular staining, samples were fixed and permeabilized using the Cytofix/Cytoperm^TM^ kit (BD Biosciences) according to manufacturers’ instructions. Antibodies used in all experiments are listed in Supplementary Table 2. Flow cytometry analysis was performed on LSR Fortessa II (BD Biosciences, BD Diva Software) and data were analyzed using FlowJo software (TreeStar, version 10.2). Cell sorting was performed using AriaII (BD Biosciences, BD Diva Software).

### Cell lines maintenance and primary cells isolation

MS5 ^53^ and engineered mesenchymal stromal cell (eMSC) lines were cultured in IMDM (Life Technologies) supplemented with 10% heat-inactivated FBS (Life Technologies), penicillin/streptomycin (Life Technologies), 50 μM β-mercaptoethanol (Life Technologies) and maintained at 37**°**C 5%CO_2_. OP9 and OP9_FLT3L were cultured in complete α-MEM (Life Technologies) supplemented with 20% not heat-inactivated FBS, penicillin/streptomycin, 50 μM β-mercaptoethanol and maintained at 37**°**C 5%CO_2_.

Cord blood units were obtained from Anthony Nolan Cell Therapy Centre (ANCTC). Peripheral blood mononuclear cells (PBMCs) were isolated by gradient centrifugation using Ficoll-Paque (GE Healthcare) and CD34^+^ hematopoietic progenitors were isolated using CD34 MicroBead Kit UltraPure (Miltenyi Biotec).

Adult peripheral blood was obtained from healthy volunteers from NHS Blood and Transplant. PBMCs were isolated by Ficoll-Paque gradient centrifugation. Cells were collected in FACS buffer and used for downstream applications.

### Human dendritic cell differentiation *in vitro*

For *in vitro* differentiation of human DCs, MS5, OP9 or eMSCs were seeded in a 96-well plate (flat bottom) at a density of 10^4^ cells/well. The following day, 10^4^ cord blood-derived CD34^+^ cells/well were seeded on top of stromal cells in complete IMDM (10% heat-inactivated FBS, penicillin/streptomycin, 50 μM β-mercaptoethanol) and maintained at 37°C 5% CO_2_. Half of the medium was replaced at day 5 and 10, and cells were collected with a solution of PBS 5 mM EDTA (at 4°C) at day 15 for flow cytometry analysis. For recombinant FLT3L experiments, human FLT3L (Celldex) 100 ng/ml was used. For transwell experiments, 24-well plates with 0.4 μm pores Transwell**^®^** inserts (Corning) were used. Stromal cells were plated at a density of 10^5^ cells/well and 7×10^4^ cord blood-derived CD34^+^ progenitors were added the following day in each well. Half of the medium was replaced in both the top and bottom well at day 5 and 10. For GM-CSF blocking experiments, 2 μg/ml human GM-CSF neutralizing antibody (R&D cat. #AF-215-SP) and isotype control (R&D cat. #AB-108-C) were added to the culture medium. The presence of human GM-CSF in the supernatant of eMSCs co-cultures with CD34^+^ HSPCs was assessed by enzyme-linked immunosorbent assay (ELISA) using the human GM-CSF ELISA MAX kit (Biolegend) as per manufacturer’s instructions.

### RNA sequencing and data processing

RNA sequencing analysis was performed in two independent experiments.

In the first experiment, human cDC1s and cDC2s from peripheral blood of n=3 healthy individuals and *in vitro* generated CD141^+^Clec9A^+^, CD14^−^CD1c^+^CD206^−^ and CD14^−^ CD1c^+^CD206^+^ cells from n=3 independent cord blood donors were FACS sorted. Up to 100 cells/subset were collected in lysis buffer (Takara Clontech) containing RNAse inhibitors. RNAseq libraries were prepared using Labcyte Echo 525 contactless liquid handling system (Labcyte Inc). In brief, ERCC mix (Thermo Fisher) was added to each sample and first strand full length cDNA was generated with a modified protocol of the SMARTseq v4 Ultra Low Input RNA Kit (Takara Clontech) using poly dT primers and a template switching oligo. Full length cDNA was amplified using SeqAmp™ DNA Polymerase (Takara Clontech). 12 ng of amplified cDNA from each sample was used to generate non-stranded RNA libraries using a modified protocol of the Ovation Ultralow System V2 1-96 kit (NuGEN). In brief, amplified cDNA was fragmented through sonication on Covaris E220 (Covaris Inc), repaired and polished followed by ligation of indexed adapters. Adapter ligated cDNA were pooled before final amplification to add flow cell primers. Libraries were sequenced on HiSeq2500 (Illumina Cambridge) for 100 cycles PE in Rapid mode. The raw sequencing data was initially processed using open source, web-based platform Galaxy (version 18.05.rc1) (https://usegalaxy.org). Reads were filtered for quality with more than 80% of the sequence having quality score > 33 using FastQC tool. Mapping against reference genome was performed with Hisat2 to the hg38 human genome. Adapter sequences were detected automatically with TrimGalore!. Reads under 20bp were discarded. All processed sequencing files were imported in Partek® Flow® software (Partek Inc., build 7.0.18.0514) and the gene count data was normalised using FPKM.

In the second experiment, human CD5^−^ cDC2, CD5^+^ cDC2, pDC and pre/AS-DC from peripheral blood of n=3 healthy individuals and cDC2, pDC and pre/AS-DC generated both *in vivo* and *in vitro* from n=2/3 independent cord blood donors were FACS sorted. Between 100 and 1000 cells/subset were collected in TRIzol^®^ (Thermo Fisher) and stored at −80°C. Frozen samples were shipped to GENEWIZ^®^ where they were processed. RNA was extracted and libraries were prepared using an ultra-low input RNA library preparation kit (Illumina). Libraries were sequenced on HiSeq2500 (Illumina).

The raw sequencing data was initially aligned on the human reference genome hg38 using STAR aligner (v2.5.3a) ^76^. Raw read counts matrix was made by STAR (with the option – *quantMode GeneCounts*).

### RNA sequencing analysis

The average gene expression of n=3 blood donors for cDC1s, n=2 blood donors for cDC2 and n=3 cord blood units for *in vitro* generated subsets were used for RNA-seq data analysis.

Hierarchical clustering was performed in Morpheus (Broad Institute, https://software.broadinstitute.org/morpheus/) using one minus Pearson’s correlation and average linkage.

Gene set enrichment analysis (GSEA) (www.broad.mit.edu/gsea) ^60^ was used to assess the expression of gene signatures specific for blood cDC1, cDC2, DC3, pDC and AS-DC in *in vitro* and *in vivo* generated subsets. To simultaneously visualize pairwise comparisons of transcriptomes from cord blood-derived cDCs, the BubbleMap module of BubbleGum ^62^ was used. Results were considered significant when the p value was below 0.05 and the FDR (false discovery rate, q) value was below 0.25. The GSEA was performed using previously published gene signatures defining blood cDC1, cDC2, DC3, pDC and AS-DC ^21^ as well as newly generated signatures using the GeneSign module of BubbleMap ^62^ (Supplementary Table 4). Briefly, the transciptome of blood cDC1s and blood CD1c^+^ cells was taken from previously published data sets ^63^. cDC1>CD1c^+^ and CD1c^+^>cDC1 signatures were defined using the “Min(test) vs Max(ref)” statistical method with a minimal linear fold change = 2 and a maximal FDR = 0.01.

Heatmaps displaying the expression pattern of gene signatures for cDC1, CD1c+ cells, pDC, AS-DC and cDC2 were generated using Morpheus (Broad Institute, https://software.broadinstitute.org/morpheus/).

The 2000 genes defining the “*in vitro* culture imprinting” were identified using Morpheus as the mean difference of expression values between two groups: the *in vitro* generated cells (including cDC1, cDC2 CD206^−^ and cDC2 CD206^+^) versus *ex vivo*-isolated subsets (blood cDC1 and blood cDC2).

The 2872 genes defining the “*in vivo* signature” were identified by DEG analysis using the R package DESeq2 (version 1.24.0) ^77^ with a Benjamin-Hochberg p-value correction ^78^ (Log2FC>1.5, adjusted p-value<0.01). The volcano plot displaying the differentially expressed genes between *in vivo* and *in vitro* differentiated cDC2 was generated using R package ggplot2 (version 3.2.1) ^79^ (Log2FC>1.5, adjusted p-value<0.05). All the analysis from the raw counts matrix were performed in Rstudio (1.2.5001) using the version 3.6.1 of R. Pathway analysis was performed using ConsensusPathDB (cpdb.molgen.mpg.de) ^80^ and WikiPathways database.

### Mass cytometry staining, barcoding, acquisition and data analysis

For mass cytometry, pre-conjugated or purified antibodies were obtained from Invitrogen, Fluidigm (pre-conjugated antibodies), Biolegend, eBioscience, Becton Dickinson or R&D Systems as listed in Supplementary Table 3. For some markers, fluorophore- or biotin-conjugated antibodies were used as primary antibodies, followed by secondary labeling with anti-fluorophore metal-conjugated antibodies (such as the anti-FITC clone FIT-22) or metal-conjugated streptavidin, produced as previously described ^81^. Briefly, 3×10^6^ cells/well in a U-bottom 96 well plate (BD Falcon) were washed once with 200 µL FACS buffer (4% FBS, 2mM EDTA, 0.05% Azide in 1X PBS) and then stained with 100 µL 200 µM cisplatin (Sigma-Aldrich) for 5 min on ice to exclude dead cells. Cells were then incubated with anti-CADM1-biotin antibody in a 50 µL reaction for 30 min on ice. Cells were washed twice with FACS buffer and incubated with 50 µL heavy-metal isotope-conjugated secondary mAb cocktail for 30 min on ice. Cells were then washed twice with FACS buffer and once with PBS before fixation with 200 µL 2% paraformaldehyde (PFA; Electron Microscopy Sciences) in PBS overnight or longer. Following fixation, the cells were pelleted and resuspended in 200 μL 1X permeabilization buffer (Biolegend) for 5 minutes at room temperature to enable intracellular labeling. Cells were then incubated with metal-conjugated anti-CD68 in a 50 µL reaction for 30 min on ice. Finally, the cells were washed once with permeabilization buffer and then with PBS before barcoding.

Bromoacetamidobenzyl-EDTA (BABE)-linked metal barcodes were prepared by dissolving BABE (Dojindo) in 100mM HEPES buffer (Gibco) to a final concentration of 2 mM. Isotopically-purified PdCl_2_ (Trace Sciences Inc.) was then added to the 2 mM BABE solution to a final concentration of 0.5 mM. Similarly, DOTA-maleimide (DM)-linked metal barcodes were prepared by dissolving DM (Macrocyclics) in L buffer (MAXPAR) to a final concentration of 1 mM. RhCl_3_ (Sigma) and isotopically-purified LnCl_3_ was then added to the DM solution at 0.5 mM final concentration. Six metal barcodes were used: BABE-Pd-102, BABE-Pd-104, BABE-Pd-106, BABE-Pd-108, BABE-Pd-110 and DM-Ln-113.

All BABE and DM-metal solution mixtures were immediately snap-frozen in liquid nitrogen and stored at −80°C. A unique dual combination of barcodes was chosen to stain each sample. Barcode Pd-102 was used at 1:4000 dilution, Pd-104 at 1:2000, Pd-106 and Pd-108 at 1:1000, Pd-110 and Ln-113 at 1:500. Cells were incubated with 100 µL barcode in PBS for 30 min on ice, washed in permeabilization buffer and then incubated in FACS buffer for 10 min on ice. Cells were then pelleted and resuspended in 100 µL nucleic acid Ir-Intercalator (MAXPAR) in 2% PFA/PBS (1:2000), at room temperature. After 20 min, cells were washed twice with FACS buffer and twice with water before a final resuspension in water. In each set, the cells were pooled from all samples, counted, and diluted to 0.5×10^6^ cells/mL. EQ Four Element Calibration Beads (DVS Science, Fluidigm) were added at a 1% concentration prior to acquisition. Cell data were acquired and analyzed using a CyTOF Mass cytometer (Fluidigm).

The CyTOF data were exported in a conventional flow-cytometry file (.fcs) format and normalized using previously-described software ^82^. Events with zero values were randomly assigned a value between 0 and –1 using a custom R script employed in a previous version of mass cytometry software ^83^. Cells for each barcode were deconvolved using the Boolean gating algorithm within FlowJo. The CD45^+^Lin (CD7/CD14/CD15/CD16/CD19/CD34)^−^ HLA-DR^+^ population of PBMC were gated using FlowJo and exported as an .fcs file. Marker expression values were transformed using the logicle transformation function ^84^. Random sub-sampling without replacement was performed to select 20,000 cell events.

The transformed values of sub-sampled cell events were then subjected to Uniform Manifold Approximation and Projection (UMAP) dimension reduction ^64, 85^ using all markers. We used the 2.4.0 release of UMAP, implemented in Python, but executed through the reticulate R package to interface R objects with Python. For both mass-cytometry datasets we used UMAP using 15 nearest neighbors (*nn*), a *min_dist* of 0.2 and euclidean distance.

Heatmaps displaying mean intensity values of CyTOF data were generated using Morpheus (Broad Institute, https://software.broadinstitute.org/morpheus/).

### Human dendritic cell differentiation *in vivo*

Human cord blood-derived CD34^+^ hematopoietic cells (5-15×10^4^ cells/plug) were injected subcutaneously along with engineered stromal cells (1:1 to 1:5 ratio HSPC/MS5) in 200 μl of ice-cold Matrigel**^®^** (BD Biosciences). Mice were sacrificed at day 12 of differentiation by cervical dislocation and Matrigel**^®^** plugs were collected. Subcutaneous Matrigel**^®^** plugs were recovered, cut in pieces and incubated in HBSS (Life Technologies) 1% FBS, 0.37 U/ml Collagenase D (Roche), 10 μg/ml DNaseI (Roche) and 1 mg/ml Dispase (Sigma-Aldrich) for 30 minutes at 37°C. After digestion, plugs were smashed on a 100 μm strainer (Corning) and cells were collected and resuspended in FACS buffer for flow cytometry analysis.

### Histology

Matrigel plugs were fixed with 1% PFA (Alfa Aesar) for 1hr at 4°C, washed and incubated in 34% sucrose solution (Sigma-Aldrich) overnight at 4°C. Plugs were embedded in Cryomatrix (Thermo Fischer) and frozen for cryostat sectioning (9 μm-thick). Sections were permeabilized using 0.5% saponin (Sigma-Aldrich), 2% BSA (Sigma-Aldrich), 1% FBS (Life Technologies) for 30 minutes at room temperature. For human DCs staining, plug sections were incubated with 1% rat anti-mouse CD16/32 (homemade) for 30 minutes to block unspecific binding sites. Sections were labeled overnight at 4°C with mouse anti-human CD1c-PE (L161, Biolegend) or mouse anti-human Clec9A-PE (8F9, Biolegend) followed by incubation for 1hr at room temperature with goat anti-mouse Cy3 (Jackson laboratory). After extensive washes, sections were labeled with mouse anti-human CD45-APC (HI30, Biolegend) for 1hr at room temperature. For human CD34^+^ progenitors staining, plugs sections were labeled overnight with purified mouse anti-human CD45 (HI30, Biolegend) followed by 1hr incubation at room temperature with goat anti-mouse Cy3. After extensive washes, sections were labeled with mouse anti-human CD34-APC (561, Biolegend) for 1hr at room temperature. To detect murine endothelial cells, sections were labeled with purified rat anti-mouse CD31 (MEC13.3, Biolegend) and mouse anti-human CD45 (HI30, Biolegend) overnight followed by 1hr incubation at room temperature with goat anti-mouse Cy3 (Jackson laboratory) and goat anti-rat Cy5 (Jackson laboratory). All sections were labeled with Hoechst (Molecular Probes, Thermo Fisher) for nuclei staining 5 minutes at room temperature and mounted with Prolong diamond (Thermo scientific). Slides were imaged using a SP5 (Leica) and analyzed with Fiji software.

### Mixed lymphocyte reaction (MLR)

Cord blood-derived DC subsets differentiated in vitro and in vivo were FACS-sorted into a V bottom 96-well plate (Corning) (10^4^ cells/well) and activated overnight (16 hours) using a TLR agonists cocktail containing LPS 10ng/ml, R848 1μg/ml and Poly(I:C) 25μg/ml.

Peripheral blood mononuclear cells (PBMC) were isolated by gradient centrifugation using Ficoll-Paque (GE Healthcare) and labeled with Cell Trace Violet (Thermo Fisher) according to manufacturer’s user guide. CTV-labeled T cells were then isolated using a Pan naïve T cell isolation kit (Miltenyi Biotec) according to manufacturer’s instructions and isolation purity (≥95%) was assessed by flow cytomery. Isolated naïve T cells (10^5^ cells/well) were seeded together with FACS-sorted DC (1:10 ratio DC/T cells) and incubated at 37°C 5% CO_2_ for 7 days.

### Statistical analysis

In all graphs each dot represents an independent cord blood donor and lines represent the median value. The number of biological as well as experimental replicates is indicated in figure legend. For each experiment, the appropriate statistical test is stated in figure legend. Statistical significance was defined as P < 0.05.

## Data availability

Data that support the findings of this study have been deposited in Gene Expression Omnibus (GEO) with the accession codes (*available before publication*).

## Acknowledgments

PG is a CNRS investigator. The research was supported by the NC3Rs, BBSRC, CRUK, Rosetree Trust and King’s Health Partners. Authors are recipient of awards and grants BBSRC-, CRUK-CIPA C57672/A22369, WWCR-18-0422, ANR-18-IDEX-0001, ANR-17-CE11-0001-01. This work was supported by the Institut Curie, Institut National de la Santé et de la Recherche Médicale, Labex DCBIOL (ANR-10-IDEX-0001-02 PSL and ANR-11-LABX0043), SIRIC INCa-DGOS-Inserm_12554, ANR JCJC (ANR-17-CE15-0004). We thank Dr Michael Ridley for assisting in RNA-seq analysis and Emily Hanton and Beth Ormrod for their technical help in performing experiments. We also want to thank Anthony Nolan Cord Blood Bank and Cell Therapy Centre for providing the cord blood units used in this study. All flow cytometry work was performed within the NIHR Biomedical Research Centre based at Guy’s and St. Thomas’ NHS Foundation Trust and King’s College London.

## Author contributions

GA, PB and OH performed the experiments. KV and YMK performed the bioinformatics analysis of RNA-seq data. CAD and EN performed CyTOF experiments and analysis. KW and AS performed RNA library preparation and sequencing. JH provided reagents and expertise for *in vitro* cultures. FG provided reagents and expertise for CyTOF analysis. GA and PG designed the experiments and wrote the manuscript. PG conceived and supervised the project.

