## Supplemental Figures for "Engineered niches support the development of human dendritic cells in humanized mice"

**Supplementary Figure 1. Stromal membrane bound FLT3L efficiently supports human cDC differentiation from CD34$^{+}$ HSPCs**

(a) Human CD123$^{+}$CD303/4$^{+}$ differentiated *in vitro* from CD34$^{+}$ cord blood-derived HSPCs cultured with MS5 expressing membrane bound FLT3L (MS5_F) or MS5 supplemented with recombinant human FLT3L (MS5+recFL) at day 15 (n=3 donors in one experiment. Line represents median; one-way ANOVA test).

(b) Flow cytometry plots and quantification of human CD123$^{+}$CD303/4$^{+}$ cells differentiated *in vitro* from cord blood-derived CD34$^{+}$ progenitors in co-culture with mouse stromal cell lines MS5 and OP9 expressing human FLT3L (MS5_F and OP9_F) at day 15 (n=4 donors in one experiment. Line represents median; one-way ANOVA test).

**Supplementary Figure 2. Stromal membrane bound SCF and CXCL12 improve the FLT3L-driven development of human cDC *in vitro***

(a) Validation of FACS-sorted stromal lines expressing single human cytokines by flow cytometry, based on the expression of fluorescent reporters as well as antibody staining of membrane-bound FLT3L and SCF. Expression of human CXCL12 was assessed by qPCR in MS5_12 after Puromycin selection.

(b) Gating strategy used to identify cord blood-derived DC subsets differentiated *in vitro*.

**Supplementary Figure 3. Stromal membrane bound SCF and CXCL12 improve the FLT3L-driven development of human cDC *in vitro***

(a) Representative FACS plots and absolute numbers of cells generated *in vitro* from $CD34^+$ HSPCs in absence of human FLT3L at day 15 (n=3 cord blood donors in 3 independent experiments. Line represents median).

(b) Absolute number of human $CD14^+CD16^-$ monocytes and $CD14^+CD16^+$ macrophages generated from $CD34^+$ HSPCs co-cultured with MS5 expressing human FLT3L (MS5_F) in combination with human SCF (S), TPO (T) and CXCL12 (12). Day15 flow cytometry analysis of n=3 cord blood donors in 3 independent experiments (line represents median; * $p<0.05$, one-way ANOVA test).

(c) Frequency of $CD206^+$ cells within the $CD14^-CD1c^+$ cells in adult peripheral blood and cord blood samples.

**Supplementary Figure 4. Human cDCs generated *in vitro* using MS5_FS12 align with circulating blood cDCs**

(a) Hierarchical clustering of primary (n=3 healthy donors) versus *in vitro* generated (n=3 cord blood donors) cDCs based on 19791 protein-coding genes.

(b) Heatmap of 2000 genes either upregulated or downregulated in *in vitro* generated cells as compared *ex vivo*-isolated blood subsets.

(c) Top 10 enriched metabolic pathways (WikiPathway) for genes upregulated or downregulated in cord blood-derived cDCs generated *in vitro* as compared to *ex vivo*-isolated subsets.

(d) GSEA of previously published gene signatures (GeneSet) of blood DC3 obtained from Villani et al., Science 2017.

(e) Surface expression of CD163 assessed by flow cytometry in *in vitro* differentiated $CD1c^{+}CD206^{-}$ and $CD1c^{+}CD206^{+}$ cells, as well as $CD14^{+}CD16^{-}$ monocytes and $CD14^{+}CD16^{+}$ macrophages in MS5_FS12 cultures.

(f) UMAP plots showing relative expression of markers detected by CyTOF in $CD45^{+}$HLA-$DR^{+}$ cells differentiated *in vitro* using MS5_FS12.

(g) Representative FACS plot and quantification of CD3+ T cells and CD19+ B cells generated in vitro using MS5_FS12.

(h) Summary of the frequency of myeloid subsets in CD45+ cells generated in vitro using MS5_FS12 (n=3-6 cord blood donors in 3 independent experiments).

(i) Absolute number of cDC1 and cDC2 generated in vitro using MS5_FS12 at day7 and day14 (n=4-7 cord blood donors in 3 independent experiments).

(l) Relative expression of TF in blood versus *in vitro*-differentiated cDC1 and cDC2.

**Supplementary Figure 5. Subcutaneous engraftment of MS5_FS12 in NSG mice results in the formation of a "synthetic niche" supporting human $CD34^+$ progenitors local maintenance and expansion**

(a) Immunofluorescence staining of plug sections displaying the interaction of $GFP^+$ MS5_FS12 (green) with human $CD45^+$ cells (red). Nuclei were stained with Hoechst (blue). Arrows show interaction of human $CD45^+$ leukocytes with $GFP^+$ MS5_FS12.

(b) Experimental strategy to assess the migration of human cells from/to MS5_FS12 plugs. CTV-labeled T cells were injected i.v. at day12 into plug-bearing mice. After 4 days, spleen and sub-cutaneous plug were recovered and the presence of human cells was assessed by flow cytometry.

**Supplementary Figure 6. The MS5_FS12 niche efficiently supports human cDC1, cDC2 and pre/AS-DCs development from CD34$^+$ HSPCs *in vivo***

(a) Flow cytometry analysis of Matrigel organoids containing either MS5_CTRL or MS5_FS12 stromal cells. Bar graphs show the absolute number of CD141$^+$Clec9A$^+$ cDC1, CD1c$^+$CD206$^-$ cDC2, CD123$^+$CD303/4$^+$ cells (n=14 donors in 6 experiments) and AXL+CD327+ pre/AS-DC (n=7 donors in 4 experiments). Line represents median; *$p<0.05$ **$p<0.01$, two-tailed paired Student t test).

(b) UMAP plots showing relative expression of markers detected by CyTOF in CD45$^+$HLA-DR$^+$ cells differentiated *in vivo* using MS5_FS12.

(c) Gating strategy used to identify cDC1, cDC2, pre/AS-DC, pDC and total CD123$^+$CD45RA$^+$ cells in two physically separated plugs containing either MS5_CTRL or MS5_FS12 injected in the same recipient. Bar graphs summarize the frequency of each subset in total CD45$^+$ cells (n=3 cord blood donors in one experiment; line represents median; * $p<0.05$, ** $p<0.01$, two-tailed paired Student t test).

(d) Gating strategy used to identify human CD15$^+$ granulocytes, CD3$^+$ T cells and CD19$^+$ B cells in vivo in MS5_FS12 organoids.

**Supplementary Figure 7. *In vivo* generated cDC2s more faithfully recapitulate the phenotype of human circulating cDC2s**

(a) Relative expression of selected markers in UMAP plots of CyTOF data comparing $CD45^{+}HLA\text{-}DR^{+}$ cells generated using MS5_FS12 stromal cells either *in vitro* or *in vivo.*

(b) Heat map of markers mean intensity (Cytof) in cDC1, cDC2 and pre/AS-DC differentiated either *in vitro* or *in vivo* using MS5_FS12 stromal cells.

(c) Heatmap of top 2000 genes either upregulated or downregulated in *in vivo* generated cells as compared *ex vivo* isolated blood subsets (top). Top 10 enriched metabolic pathways (WikiPathway) for genes upregulated or downregulated in cord blood-derived cDCs generated *in vivo* as compared to *ex vivo* isolated subsets (bottom).

(d) Top 10 enriched metabolic pathways (WikiPathway) for genes upregulated in cord blood-derived cDC2 generated *in vitro* (left). Top 10 enriched metabolic pathways (WikiPathway) for genes upregulated in cord blood-derived cDC2 generated *in vivo* (right).

**Supplementary Figure 8. cDC2 generated *in vitro* and *in vivo* recapitulate functional features ascribed to blood cDC2.**

(a) Gating strategy used to FACS-sort DC2, pDC and pre/AS-DC differentiated both in vitro and in vivo using MS5_FS12 stromal niche.

(b) Representative FACS plots of mixed lymphocyte reaction (MLR) using *ex vivo* isolated primary cDC2 and $CD14^{+}$ monocytes. FACS-sorted DC subsets were activated overnight (16 hours) using a TLR agonist cocktail (LPS 10ng/ml, R848 1µg/ml and Poly(I:C) 25µg/ml) and co-cultured with CTV-labeled naïve T cells for 5 days.

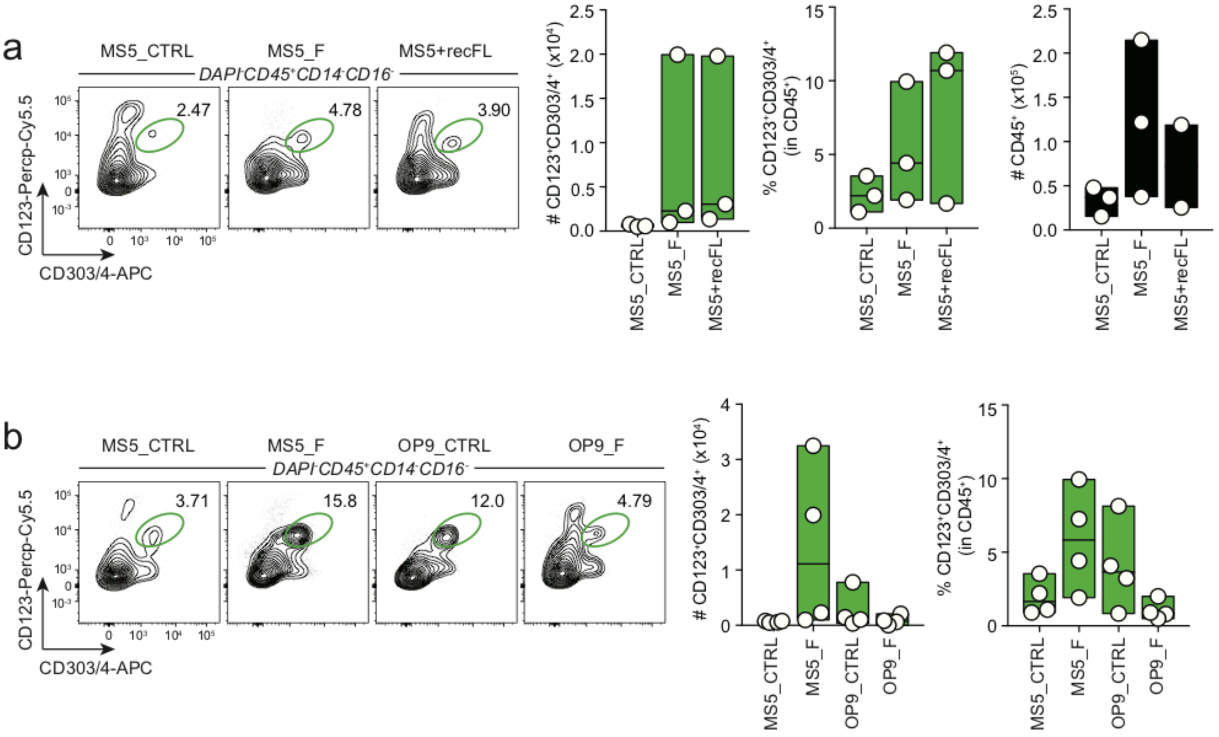
a
MS5_CTRL
MS5_F
MS5+recFL
DAPI⁻CD45⁺CD14⁻CD16⁻
2.47
4.78
3.90
CD123-Percp-Cy5.5
CD303/4-APC
# CD123⁺CD303/4⁺ (x10⁴)
% CD123⁺CD303/4⁺ (in CD45⁺)
# CD45⁺ (x10⁵)
b
MS5_CTRL
MS5_F
OP9_CTRL
OP9_F
DAPI⁻CD45⁺CD14⁻CD16⁻
3.71
15.8
12.0
4.79
CD123-Percp-Cy5.5
CD303/4-APC
# CD123⁺CD303/4⁺ (x10⁴)
% CD123⁺CD303/4⁺ (in CD45⁺)

### Supplementary Fig.2

a

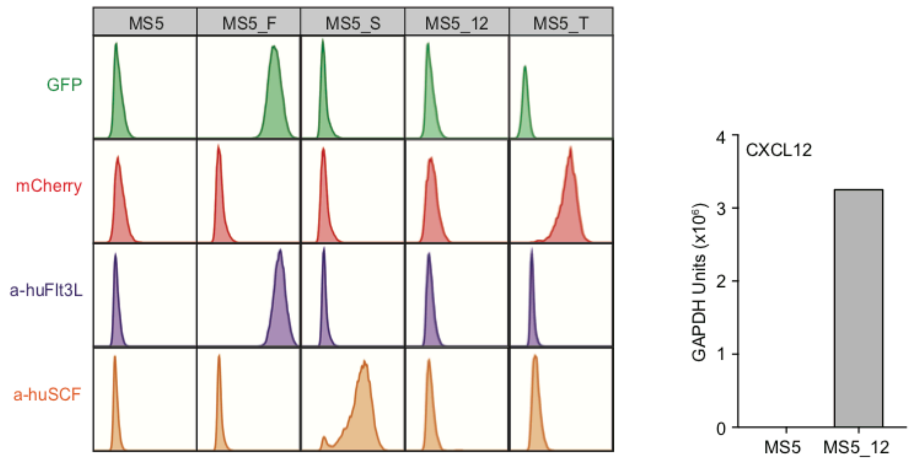


| | CTRL | F | S | T | 12 | FS | FT | F12 | ST | S12 | T12 | FST | FT12 | FS12 | ST12 | FST12 |
|---|---|---|---|---|---|---|---|---|---|---|---|---|---|---|---|---|
| huFlt3L-IRES-GFP | - | + | - | - | - | + | + | + | - | - | - | + | + | + | - | + |
| huSCF (no tag) | - | - | + | - | - | + | - | - | + | + | - | + | - | + | + | + |
| huTPO-IRES-mCherry | - | - | - | + | - | - | + | - | + | - | + | + | + | - | + | + |
| huCXCL12-IRES-PuroR | - | - | - | - | + | - | - | + | - | + | + | - | + | + | + | + |

b

Gating strategy engineered MS5 screening:

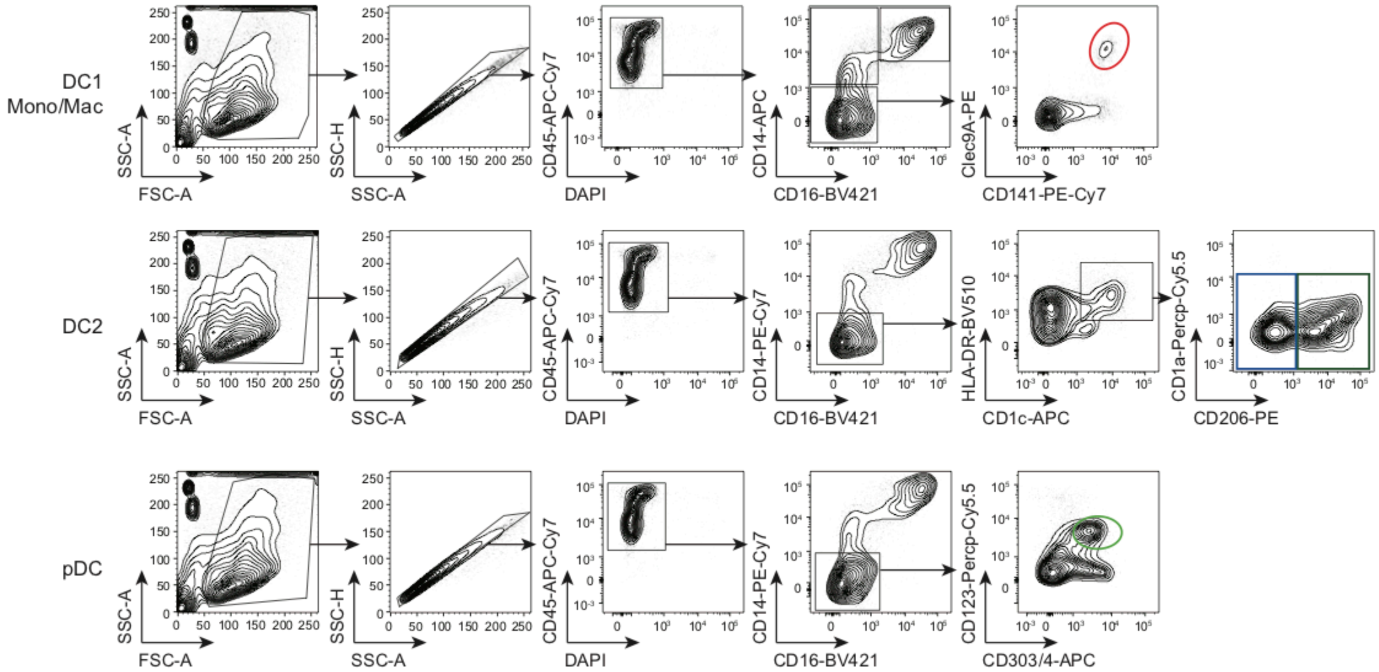

Supplementary Fig.3

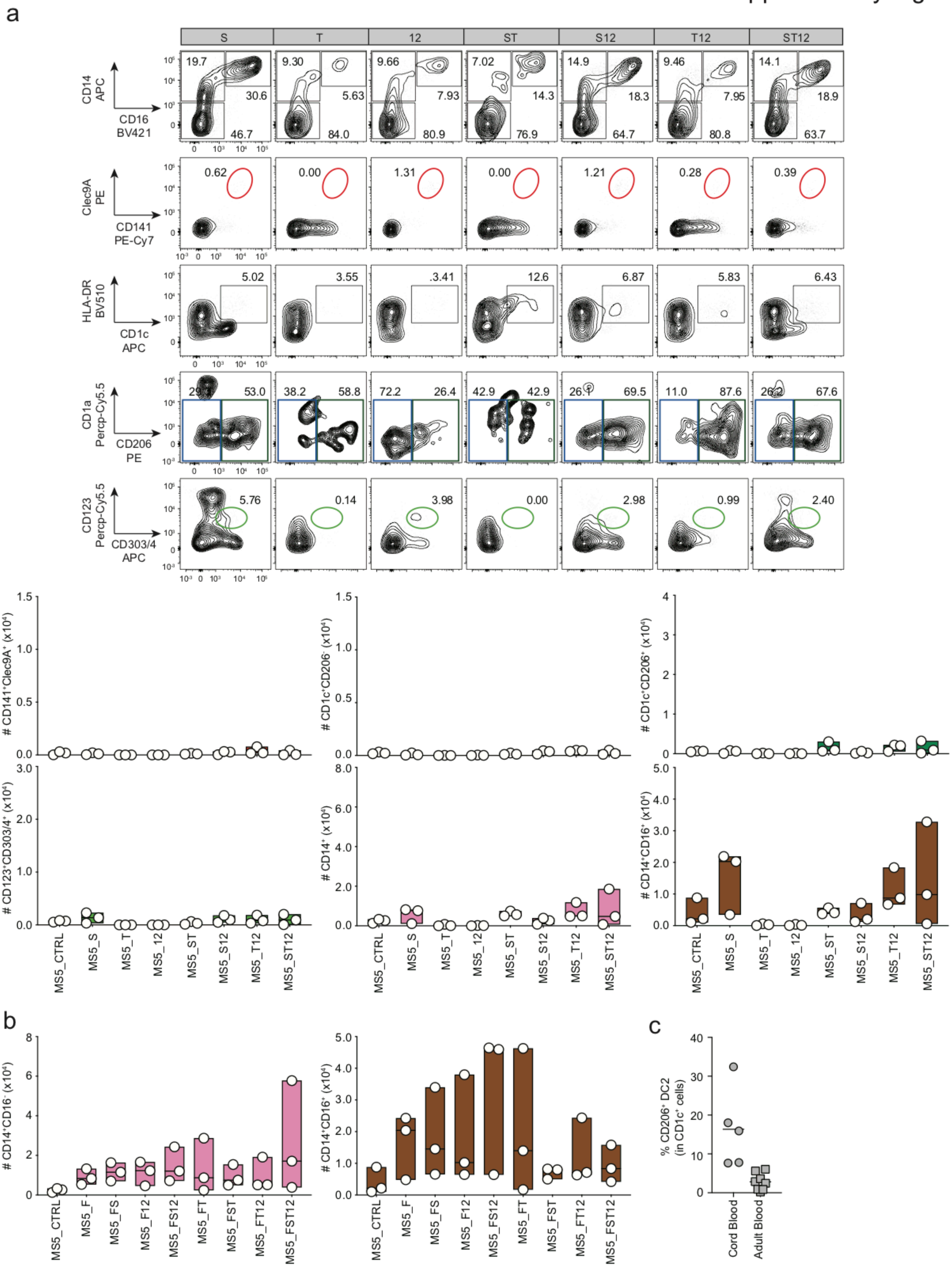

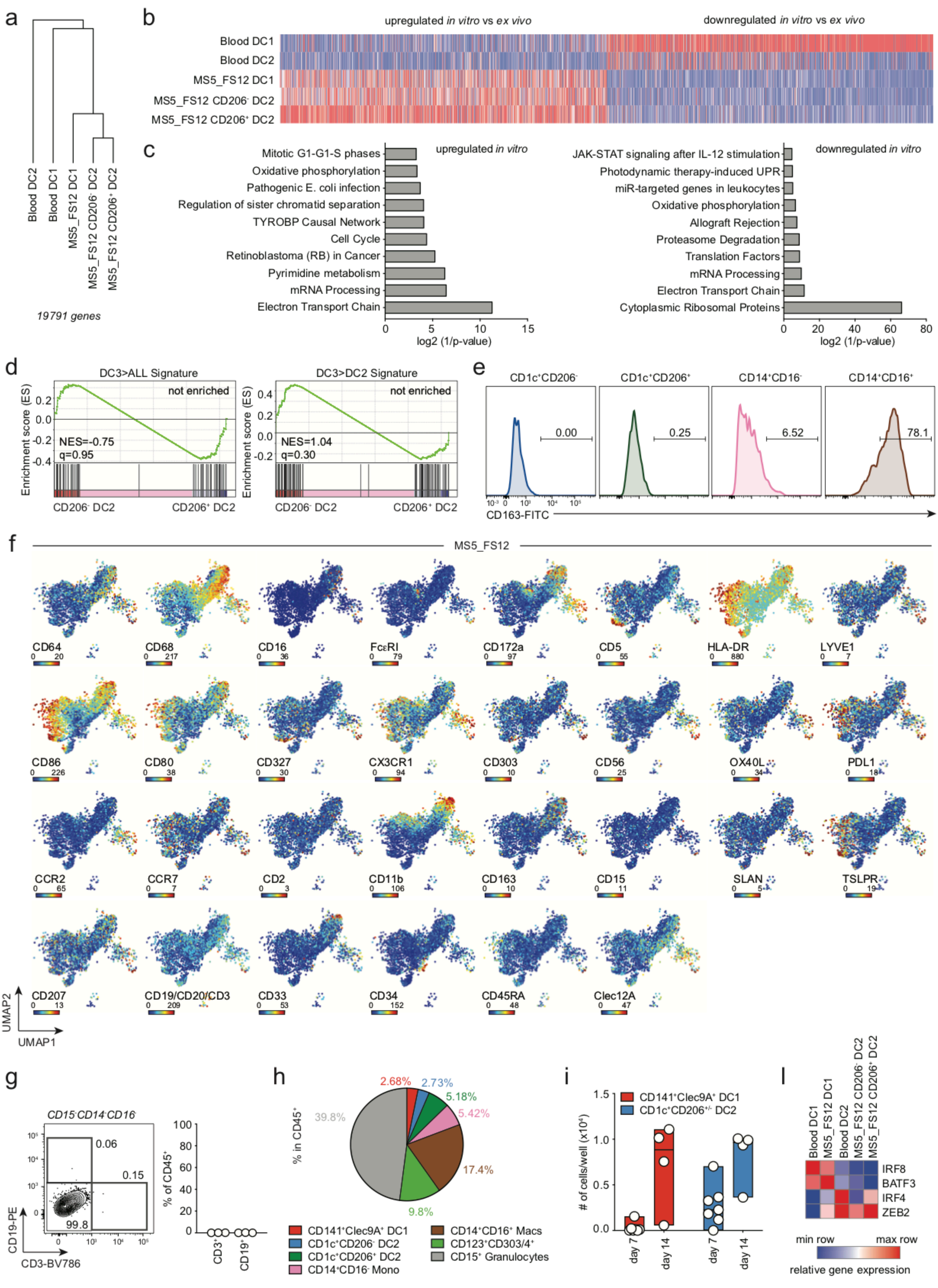
a
b
upregulated in vitro vs ex vivo
downregulated in vitro vs ex vivo
Blood DC1
Blood DC2
MS5_FS12 DC1
MS5_FS12 CD206- DC2
MS5_FS12 CD206+ DC2
19791 genes
c
upregulated in vitro
Mitotic G1-G1-S phases
Oxidative phosphorylation
Pathogenic E. coli infection
Regulation of sister chromatid separation
TYROBP Causal Network
Cell Cycle
Retinoblastoma (RB) in Cancer
Pyrimidine metabolism
mRNA Processing
Electron Transport Chain
log2 (1/p-value)
downregulated in vitro
JAK-STAT signaling after IL-12 stimulation
Photodynamic therapy-induced UPR
miR-targeted genes in leukocytes
Oxidative phosphorylation
Allograft Rejection
Proteasome Degradation
Translation Factors
mRNA Processing
Electron Transport Chain
Cytoplasmic Ribosomal Proteins
log2 (1/p-value)
d
DC3>ALL Signature
not enriched
NES=-0.75
q=0.95
DC3>DC2 Signature
not enriched
NES=1.04
q=0.30
Enrichment score (ES)
CD206- DC2
CD206+ DC2
e
CD1c+CD206-
CD1c+CD206+
CD14+CD16-
CD14+CD16+
0.00
0.25
6.52
78.1
CD163-FITC
f
MS5_FS12
CD64
CD68
CD16
FcεRI
CD172a
CD5
HLA-DR
LYVE1
CD86
CD80
CD327
CX3CR1
CD303
CD56
OX40L
PDL1
CCR2
CCR7
CD2
CD11b
CD163
CD15
SLAN
TSLPR
CD207
CD19/CD20/CD3
CD33
CD34
CD45RA
Clec12A
UMAP1
UMAP2
g
CD15-CD14-CD16-
0.06
0.15
99.8
CD19-PE
CD3-BV786
% of CD45+
CD3+
CD19+
h
% in CD45+
2.68%
2.73%
5.18%
5.42%
39.8%
17.4%
9.8%
CD141+Clec9A+ DC1
CD1c+CD206- DC2
CD1c+CD206+ DC2
CD14+CD16- Mono
CD14+CD16+ Macs
CD123+CD303/4+
CD15+ Granulocytes
i
CD141+Clec9A+ DC1
CD1c+CD206+/- DC2
### of cells/well (x10^4)
day 7
day 14
l
Blood DC1
MS5_FS12 DC1
Blood DC2
MS5_FS12 CD206- DC2
MS5_FS12 CD206+ DC2
IRF8
BATF3
IRF4
ZEB2
min row
max row
relative gene expression

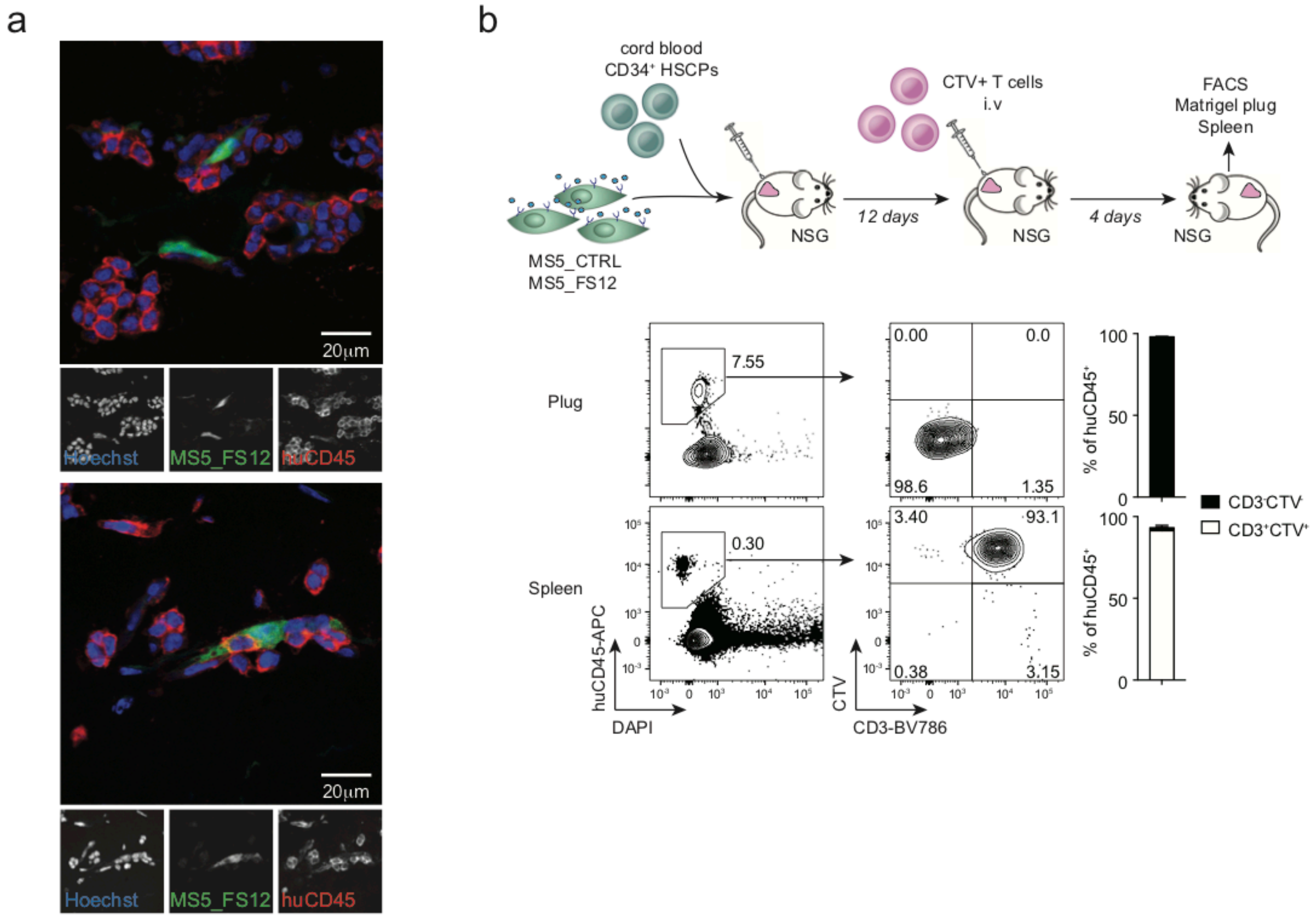
a
20µm
Hoechst
MS5_FS12
huCD45
20µm
Hoechst
MS5_FS12
huCD45
b
cord blood
CD34+ HSCPs
MS5_CTRL
MS5_FS12
NSG
12 days
CTV+ T cells
i.v
NSG
4 days
FACS
Matrigel plug
Spleen
NSG
Plug
7.55
0.00
0.0
98.6
1.35
Spleen
0.30
3.40
93.1
0.38
3.15
huCD45-APC
DAPI
CTV
CD3-BV786
% of huCD45+
CD3-CTV-
CD3+CTV+

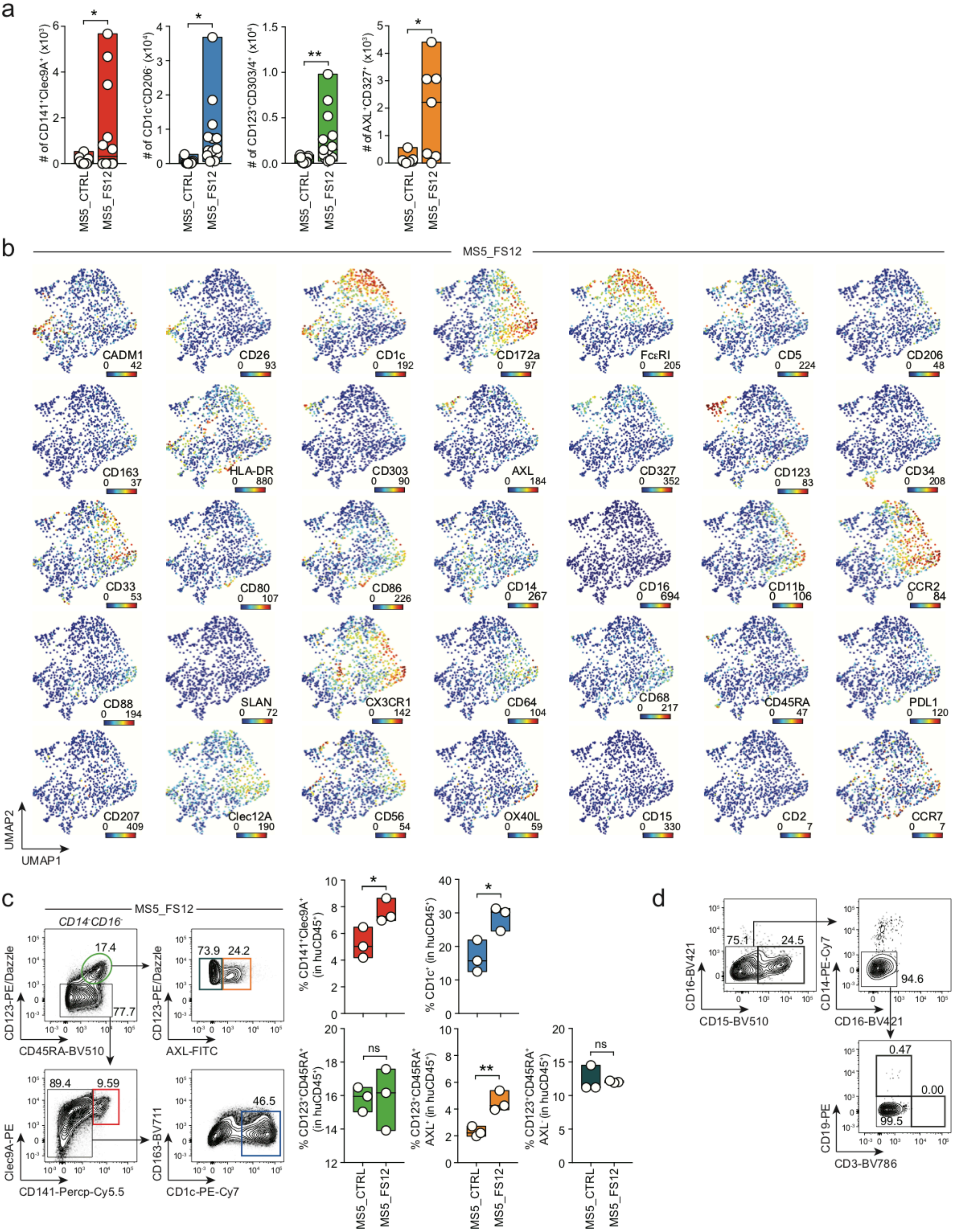
a
### of CD141+Clec9A+ (x10³)
# of CD1c+CD206- (x10⁴)
# of CD123+CD303/4+ (x10⁴)
### of AXL+CD327+ (x10³)
MS5_CTRL
MS5_FS12
b
MS5_FS12
CADM1
CD26
CD1c
CD172a
FcεRI
CD5
CD206
CD163
HLA-DR
CD303
AXL
CD327
CD123
CD34
CD33
CD80
CD86
CD14
CD16
CD11b
CCR2
CD88
SLAN
CX3CR1
CD64
CD68
CD45RA
PDL1
CD207
Clec12A
CD56
OX40L
CD15
CD2
CCR7
UMAP2
UMAP1
c
MS5_FS12
CD14-CD16-
CD123-PE/Dazzle
CD45RA-BV510
AXL-FITC
Clec9A-PE
CD141-Percp-Cy5.5
CD163-BV711
CD1c-PE-Cy7
% CD141+Clec9A+ (in huCD45+)
% CD1c+ (in huCD45+)
% CD123+CD45RA+ (in huCD45+)
% CD123+CD45RA+ AXL+ (in huCD45+)
% CD123+CD45RA+ AXL- (in huCD45+)
ns
d
CD16-BV421
CD15-BV510
CD14-PE-Cy7
CD19-PE
CD3-BV786

Supplementary Fig.7

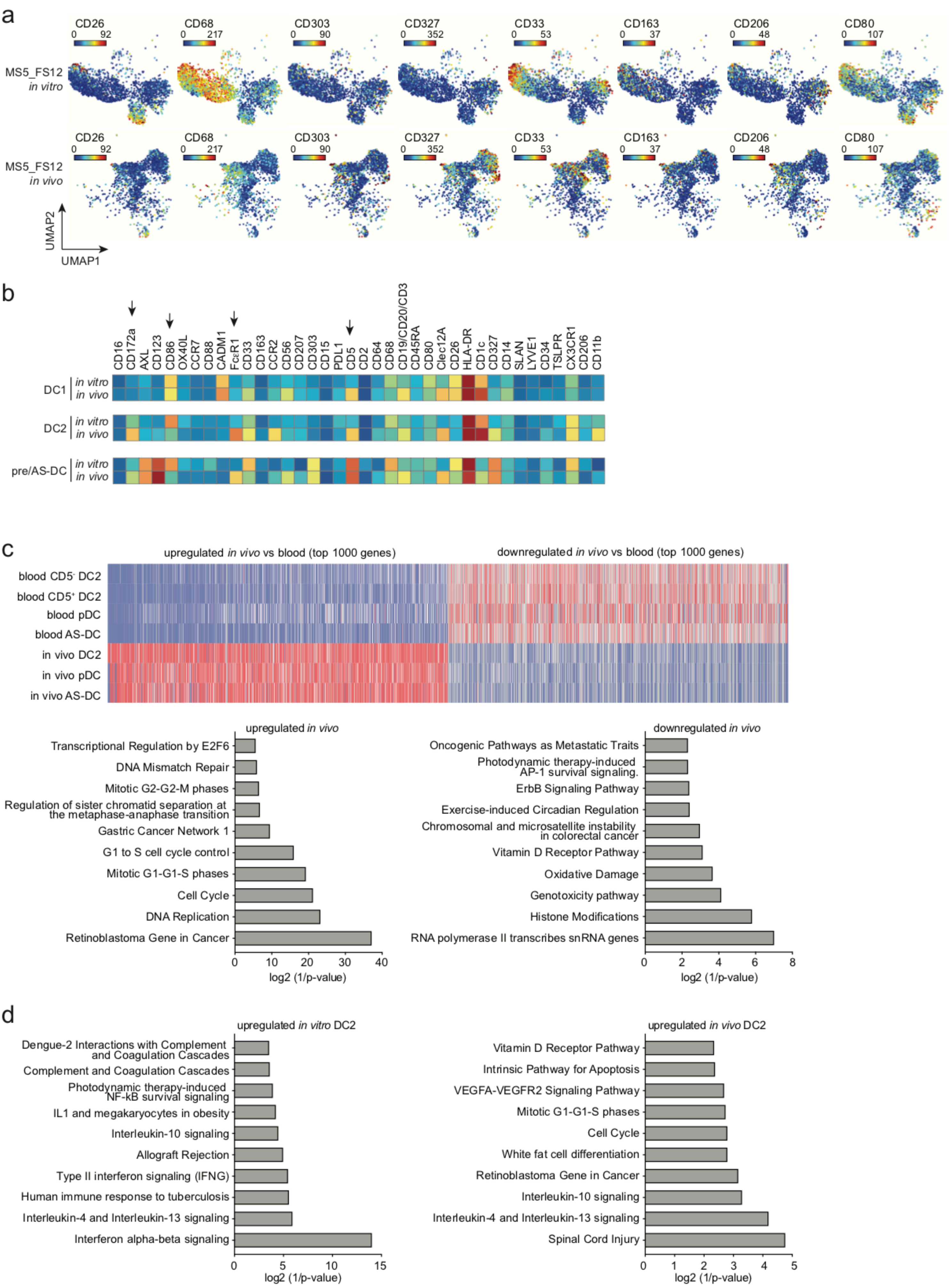

Supplementary Fig.8

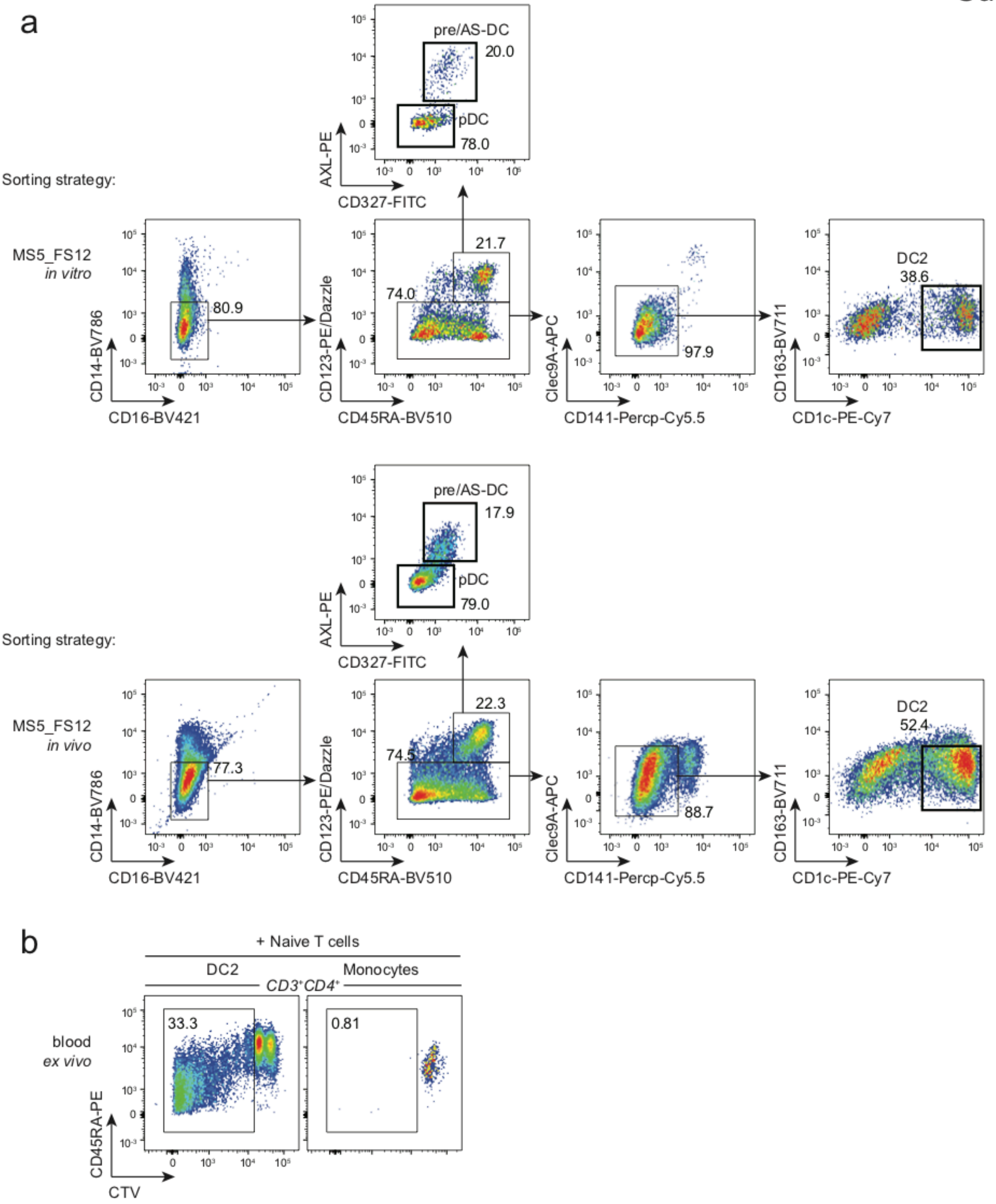
